# The PSI–NDH supercomplex prevents chilling-induced PSI photoinhibition

**DOI:** 10.64898/2026.05.11.724080

**Authors:** Ko Takeuchi, Shintaro Harimoto, Kentaro Ifuku

**Author notes:** Corresponding author: Kentaro Ifuku. These authors contributed equally to this work (Ko Takeuchi, Shintaro Harimoto).

## Abstract

Chilling stress induces photosystem I (PSI) photoinhibition in chilling-sensitive cucumber, in which insufficient activity of the chloroplast NADH dehydrogenase-like complex (NDH) leads to PSI over-reduction and damage. However, it is not yet clear whether these findings can be generalized to other species or what the molecular mechanism underlying impaired NDH function is. In this study, we first examined whether NDH is essential for PSI protection under chilling stress using an NDH-deficient rice mutant. Compared with wild-type plants, the NDH-deficient mutant exhibited enhanced PSI over-reduction and pronounced PSI photoinhibition under chilling stress. In contrast, rice plants expressing flavodiiron protein (FLV), which functions as an alternative electron acceptor downstream of PSI, did not exhibit PSI photoinhibition under chilling stress, demonstrating that electron sink capacity of NDH is important for PSI protection under chilling stress. Furthermore, analysis of the factors responsible for NDH dysfunction under chilling stress in cucumber revealed that chilling stress destabilizes the PSI–NDH supercomplex, leading to NDH monomerization and a consequent loss of NDH activity. This NDH monomerization is likely attributable to chilling-induced damage to the light-harvesting complex Lhca, which mediates the association between PSI and NDH. Together, these results indicate that NDH is essential for protecting PSI from photoinhibition under chilling stress in both rice and cucumber, and that chilling-induced destabilization of the PSI–NDH supercomplex represents a key molecular mechanism underlying PSI over-reduction and photoinhibition.

## Introduction

Chilling stress reduces CO_2_ fixation capacity and induces photoinhibition of photosystems (PS), thereby limiting crop productivity (Kingston-Smith *et al*., 1997; Allen and Ort, 2001; Zhou *et al*., 2007). Under chilling stress, plants are at risk of both PSII and PSI photoinhibition (Terashima *et al*., 1994; Nishiyama *et al*., 2001; Tyystjärvi, 2008). PSII photoinhibition is often reversible due to its efficient repair and has little impact on plant growth under short-term chilling stress. Rather, PSII photoinhibition can function as a survival strategy that helps prevent PSI photoinhibition (Ivanov *et al*., 1998; Kim *et al*., 2005; Tikkanen *et al*., 2014; Zhang *et al*., 2016; Lima-Melo *et al*., 2019; Obara *et al*., 2022; Messant *et al*., 2024; Napaumpaiporn *et al*., 2024; Ozawa *et al*., 2024; Takeuchi *et al*., 2025*b*). In contrast, PSI is repaired much more slowly, and once photoinhibited, causes severe leaf damage (Kudoh and Sonoike, 2002; Zivcak *et al*., 2015; Amin *et al*., 2024). In several plant species, including cucumber, tomato, potato, cotton, and sugarcane, PSI photoinhibition is readily induced under chilling stress, resulting in substantial growth inhibition (Havaux and Davaud, 1994; Erling Tjus *et al*., 1998, 1999; Kornyeyev *et al*., 2003; Govindachary *et al*., 2004; Chen *et al*., 2024; Li *et al*., 2024; Takeuchi *et al*., 2024; Zhang Y *et al*., 2024).

PSI photoinhibition in intact leaves has been investigated for several decades, with the first report dating back to studies in cucumber in 1994 (Sonoike and Terashima, 1994; Terashima *et al*., 1994). Subsequently, PSI photoinhibition in intact leaves has also been reported in plant species other than cucumber under chilling stress, as well as under non-chilling conditions such as fluctuating light and low CO_2_. (e.g. Ivanov *et al*., 1998; Takagi *et al*., 2017*b*, 2019; Huang *et al*., 2019; Wada *et al*., 2019; Terashima *et al*., 2021; Sun *et al*., 2023; Takagi and Tani, 2023; Grebe *et al*., 2024; Tiwari *et al*., 2024). The primary cause of PSI photoinhibition is oxidative damage resulting from over-reduction of PSI. In PSI, the redox potentials of electron carriers (phylloquinone A_1_ and iron-sulfur clusters F_X_, F_A_/F_B_) are sufficiently low to reduce oxygen to superoxide (O_2_^•–^) (Kozuleva and Ivanov, 2010; Cherepanov *et al*., 2017; Kozuleva *et al*., 2021; Degen and Johnson, 2024). In addition, when the acceptor side of PSI becomes fully reduced, charge recombination between radical pairs P700^+^/A_0_^−^ or P700^+^/A^1–^ can generate the triplet state of P700 (^3^P700), which produces singlet oxygen (^1^O_2_) (Shuvalov *et al*., 1986; Kim *et al*., 2001; Rajagopal *et al*., 2005; Rutherford *et al*., 2012; Shimakawa *et al*., 2024).

To protect chloroplasts from this oxidative damage, regulation of electron transport on the PSI donor side is critical, particularly control of electron flow at the cytochrome *b*_6_*f* complex (photosynthetic control) and suppression of electron transport through PSII photoinhibition (Foyer *et al*., 1990; Takagi *et al*., 2017*a*; Miyake, 2020; Degen and Johnson, 2024; Takeuchi *et al*., 2025*b*). In parallel, alternative electron sinks on the PSI acceptor side are also essential for preventing PSI photoinhibition (Shimakawa and Miyake, 2018; Luu Trinh *et al*., 2021; Rodriguez-Heredia *et al*., 2022). In plants other than angiosperms, flavodiiron protein (FLV) plays a major role as downstream electron sinks of PSI. FLV accept electrons from PSI and reduce oxygen to water in a safe manner, thereby contributing to the suppression of PSI photoinhibition (Helman *et al*., 2003; Shimakawa *et al*., 2019). However, most terrestrial plants have lost FLV during evolution and are therefore thought to compensate through enhanced photosynthetic control and the utilization of alternative pathways such as photorespiration and cyclic electron flow (CEF) (Hanawa *et al*., 2017; Hani *et al*., 2024; Satoh *et al*., 2025).

The chloroplast NADH dehydrogenase-like complex (NDH) has been proposed to suppress oxidative damage to PSI by transferring electrons from ferredoxin (Fd) to plastoquinone (PQ) (Shikanai, 2016, 2024; Richardson *et al*., 2021; Shikanai *et al*., 2025). NDH is composed of five subcomplexes: SubM (NdhA–NdhG), SubA (NdhH–NdhO), SubE (NdhS–NdhV), SubB (PnsB1–PnsB5) and the luminal subcomplex SubL (PnsL1–PnsL5) (Ifuku *et al*., 2011). NDH is sandwiched between two PSI complexes to form the PSI–NDH supercomplex (sc) (Kouřil *et al*., 2014; Schuller *et al*., 2019; Pan *et al*., 2020; Shen *et al*., 2022; Su *et al*., 2022; Introini *et al*., 2025). The association between PSI and NDH is mediated by the specialized light-harvesting proteins Lhca5 and Lhca6 (Otani *et al*., 2017, 2018; Kato *et al*., 2018*b*, 2021). In this study, PSI associated with NDH via Lhca5 is referred to as right PSI, whereas PSI associated via Lhca6 is referred to as left PSI (Su *et al*., 2022). In some cases, additional PSI complexes associate with the left PSI to form a larger PSI–NDH sc (Yadav *et al*., 2017). Despite these extensive structural studies, NDH-deficient mutants of *A. thaliana* exhibit phenotypes comparable to those of wild-type plants (Shikanai, 2016), leaving its role in environmental stress tolerance in C₃ plants largely unresolved.

The importance of NDH under chilling stress has been suggested by observations that NDH accumulation increases at low temperature and that reduced NDH levels in maize decrease tolerance to chilling stress (Bernhard Teicher *et al*., 2000; Urzinger *et al*., 2025). More recently, we demonstrated that chilling-induced PSI photoinhibition in cucumber is caused not only by impairment of chloroplast ATPase (Peeler and Naylor, 1988; Terashima *et al*., 1989, 1991*a*,*b*), but also by a decline in NDH activity (Takeuchi *et al*., 2022, 2025*a*). In chilling-sensitive cucumber cultivars, NDH activity was lost within 1 h of chilling stress, leading to accumulation of reduced ferredoxin and subsequent PSI photoinhibition (Takeuchi *et al*., 2025*a*). In contrast, in chilling-tolerant cucumber cultivars, NDH functioned normally under chilling stress, and NDH contributed to suppression of PSI photoinhibition by dissipating excess electrons downstream of PSI (Takeuchi *et al*., 2025*a*).

In this study, we addressed two questions to clarify the role of NDH in chilling-induced PSI photoinhibition. First, we analyzed chilling-induced PSI photoinhibition in an NDH-deficient rice mutant to examine whether NDH is indispensable for PSI protection under chilling stress in plants more generally, rather than only in cucumber. Second, we investigated the factors responsible for NDH inactivation under chilling stress using cucumber as a model, in which NDH inactivation and PSI photoinhibition are particularly pronounced.

## Materials and Methods

### Plant materials, growth conditions, and chilling stress treatments

Seeds of japonica rice (*Oryza sativa* L.) cultivar Hitomebore or Nipponbare (WT), the NDH-deficient mutant *crr6* (background: Hitomebore), and FLV overexpression line (background: Nipponbare), were soaked for 2 days in distilled water containing a fungicide (Trifumin; Nippon Soda, Tokyo, Japan). The seeds were transferred to a hydroponic medium and grown in a growth chamber under a 16 h light (250–300 µmol photons m^-2^ s^-1^) / 8 h dark photoperiod at 27 °C for 1 month. The hydroponic medium consisted of the following components in tap water (Takeuchi et al., 2024): 0.5 mM CaCl_2_, 0.25 mM KH_2_PO_4_, 1 mM (NH_4_)_2_SO_4_, 0.5 mM MgCl_2_, 0.5 mM KCl, 2.86 mg L^-1^ H_3_BO_3_, 1.81 mg L^-1^ MnCl_2_·4H_2_O, 0.08 mg L^-1^ CuSO_4_·5H_2_O, 0.22 mg L^-1^ ZnSO_4_·7H_2_O, 0.1 mg L^-1^ (NH_4_)_6_Mo_7_O_24_·4H_2_O, and 90 µM Fe(III)-EDTA. The medium was refreshed twice per week. Seeds of cucumber (*Cucumis sativus* L. ‘High Green 21’; HG) were purchased from Saitama Genshu Ikuseikai (Saitama, Japan). Plants were grown in plastic pots containing soil in a growth chamber under a 16 h light (250–300 µmol photons m^-2^ s^-1^) / 8 h dark photoperiod at 27 °C. Fully expanded leaves from 2–3-week-old plants were used for all analyses. For chilling stress treatment, plants were transferred to a cold room at 4 °C and exposed to light (180–250 µmol photons m^-2^ s^-1^). Plants not subjected to chilling stress were used as controls.

### Measurement of Chl fluorescence and P700 redox state

Chl fluorescence and redox changes of P700 were measured using a Dual-PAM 100 measuring system (Heinz Walz GmbH, Effeltrich, Germany) at 23 °C immediately after chilling stress, as previously described (Takeuchi *et al*., 2022, 2025*a*). The maximum quantum yield of PSII in the dark was calculated as F_v_/F_m_. The effective quantum yield of PSII was calculated as Y(II) (ΦII) = (F_m_’−F_s_)/F_m_’. The quantum yields of nonregulated and regulated energy dissipation in PSII were calculated as Y(NO) (ΦNO) = F_s_/F_m_, Y(NPQ) (ΦNPQ) = F_s_/F_m_’−F_s_/F_m_, and NPQ = F_m_/F_m_’−1, respectively. The F_o_ and F_m_ levels, representing the minimum and maximum fluorescence, respectively, were measured after leaves were kept in the dark for at least 20 min. Chilled plants were kept in the dark in a cold room (4 °C), whereas control plants were kept at 23 °C. After the onset of AL illumination, saturation pulses (SP; 300 ms and 20,000 µmol photons m^-2^ s^-1^) were applied to monitor F_m_’ (maximum fluorescence under light) and F_s_ (steady-state fluorescence under light). Redox changes in P700 were analyzed by monitoring the absorbance changes at 830 and 875 nm (Klughammer and Schreiber, 2008). The ratio of P700^+^ under AL was calculated as Y(ND) = P/P_m_. The ratio of P700 that cannot be oxidized by SP was calculated as Y(NA) = (P_m_−P_m_’)/P_m_. Y(I) was calculated as (P_m_’−P)/P_m_ (Klughammer and Schreiber, 1994). P_m_ was determined by applying SP after FR light illumination. P_m_’ represents the maximum level of P700^+^ under AL+SP, and P represents the steady-state P700^+^ level recorded immediately before the application of SP. It should be noted that Y(I) depends on the rate-limiting step of the P700 redox cycle during SP and does not exactly reflect the quantum yield of PSI (Furutani *et al*., 2022; Grebe *et al*., 2024).

### Measurement of NDH activity

NDH activity was assessed by monitoring the transient increase in Chl fluorescence after AL was turned off, referred to as the post-illumination fluorescence rise (PIFR), using MINI-PAM (Heinz Walz GmbH) or Dual PAM 100 at 23 °C immediately after chilling stress. Leaves were kept in the dark for at least 20 min before exposure to AL (60 µmol photons m^-2^ s^-1^) for 4–5 min. The subsequent transient increase in Chl fluorescence was monitored under the measuring light immediately after AL was turned off.

### Isolation of thylakoid membranes

Fresh leaves collected before and immediately after 1 h of chilling stress were cut into small strips and homogenized in 50 mL of ice-cold HEPES buffer (0.3 M sorbitol, 1 mM MgCl_2_, 1 mM MnCl_2_, 2 mM EDTA-Na_2_, 30 mM KCl, 0.25 mM KH_2_PO_4_, and 50 mM HEPES-KOH (pH 7.6)) containing 0.1% (w/v) bovine serum albumin using an ACE homogenizer AM-8 (Nissei, Aichi, Japan) at maximum voltage for 10 s, repeated three times. The homogenate was filtered through two layers of Miracloth (EMD Millipore, Danvers, MA, USA) and centrifuged at 1,030×g for 5 min at 4 °C. The pellet was resuspended in 1 mL of HEPES buffer and loaded into a tube containing an 80% (w/v) Percoll in HEPES buffer. The tubes were centrifuged at 1,030×g at 4 °C for 20 min, and the green layer on the top of the 80% Percoll was collected as the thylakoid fraction. To remove Percoll, the thylakoid fraction was diluted with three volumes of washing buffer containing 1 mM MgCl_2_, 1 mM MnCl_2_, 2 mM EDTA-Na_2_, 30 mM KCl, 0.25 mM KH_2_PO_4_, and 50 mM HEPES-KOH (pH 7.6), followed by centrifugation at 1,030×g for 5 min at 4 °C.

### *In vitro* Fd-dependent PQ reduction assay

The Fd-dependent PQ reduction assay was performed as previously described (Endo *et al*., 1998; Okegawa and Motohashi, 2020). Isolated thylakoids (20 µg Chl mL^−1^) collected before and after 1 h of chilling stress were suspended in rupture buffer containing 15 mM MgCl_2_, 1 mM MnCl_2_, 2 mM EDTA-Na_2_, 30 mM KCl, 0.25 mM KH_2_PO_4_, and 50 mM HEPES-KOH (pH 7.6). Fd-dependent PQ reduction activity was assayed by monitoring the increase in Chl fluorescence following the addition of 250 µM NADPH and 2.5 µM spinach Fd under weak measuring light using MINI PAM. Antimycin A (AA) was added to a final concentration of 5 µM. In this analysis, the effects of AA on PSII were considered negligible. Chl fluorescence was normalized by F_o_ and F_m_.

### Blue-native PAGE (BN-PAGE), SDS-PAGE, and immunoblot analysis

BN-PAGE was performed as previously described (Järvi *et al*., 2011). Thylakoid membranes were solubilized with n-dodecyl-β-D-maltoside (β-DM) on ice for 1 min, and the insoluble fraction was removed by centrifugation at 21,500 × g for 2 min. The resulting supernatant containing 5 µg Chl was loaded onto a Native-PAGE 4–13% gradient gel. Electrophoresis was performed using the myPowerII 300 system AE-8135 (ATTO, Tokyo, Japan) at 200 V and 1 mA for 13 h at 4 °C. For two-dimensional (2D)-BN/SDS-PAGE analysis, BN-PAGE gel lanes were excised and soaked in 1% SDS solubilization buffer containing 50 mM dithiothreitol for 30 min at 23 °C. Each lane containing denatured proteins was placed on top of a 1.0-mm-thick 12% wide-range gel and electrophoresed at 40 mA. The gels were subsequently subjected to immunoblotting. SDS-PAGE and immunoblot analyses were performed as described previously (Che *et al*., 2020). Thylakoid membrane protein samples were loaded onto 12% wide-range gels and separated by SDS-PAGE. After electrophoresis, the gels were desalted and equilibrated in transfer buffer containing 25 mM Tris, 192 mM glycine, 20% (v/v) methanol, 0.03% (w/v) SDS, and proteins were transferred to 0.45 µm polyvinylidene difluoride (PVDF) membranes. The membranes were blocked with a PVDF blocking reagent (TOYOBO, Osaka, Japan) and subsequently incubated with specific antibodies. Detection was performed using Amersham ECL Prime Western Blotting Detection Reagent (Cytiva, Tokyo, Japan) and imaged with the ChemiDoc Touch Imaging System (Bio-Rad, Hercules, CA, USA). Anti-NdhH, PnsB1, PnsL1, NdhT, PsaA, Lhca2, Lhca3, Lhca5, and Cyt *f* specific antibodies were purchased from Agrisera, and anti-Lhca6 specific antibody was purchased from PhytoAB.

### Statistical analysis

Statistical analyses performed using Student’s *t*-test. ALL calculations were based on at least three independent biological replicates.

## Results

### Chilling-induced photoinhibition in an NDH-deficient rice mutant

NDH accepts electrons from Fd and contributes to maintaining PSI in an oxidized state. Previous studies have shown that NDH inactivation under chilling stress induces PSI photoinhibition in cucumber (Takeuchi *et al*., 2025*a*). To address whether the role of NDH in protecting PSI under chilling stress extends beyond cucumber, and how NDH deficiency affects the susceptibility of PSI to photoinhibition under chilling stress, we analyzed chilling-induced PSI photoinhibition in an NDH-deficient rice mutant. The NDH-deficient mutant *crr6* (cv. Jp Hitomebore), in which SubA of the PSI–NDH sc fails to accumulate (Yamori *et al*., 2011), lacked NDH activity (Supplementary Fig. S1). Rice plants were grown at 27 °C for one month and then subjected to chilling stress for 24 h (4 °C, under light). Changes in photosynthetic activity were monitored during chilling stress and after 24 h of recovery (Fig. 1A). Notably, the total amount of active PSI (Pm), calculated from the maximum photo-oxidizable P700, decreased markedly in *crr6* under chilling stress (Fig. 1B). After 6 h of chilling stress, no decrease in active PSI amount was observed in the wild type (WT), whereas it declined to approximately 0.8 in *crr6*. After 24 h of chilling stress, the active PSI amount in WT decreased to approximately 0.8, indicating that chilling-induced PSI photoinhibition also progressed in rice WT. In particular, *crr6* showed a much larger decrease, with the active PSI amount declining to approximately 0.4, demonstrating pronounced PSI photoinhibition. The maximum quantum yield of PSII (Fv/Fm) decreased to approximately 0.6 after 6 h of chilling stress in both WT and *crr6* and further declined to approximately 0.2–0.3 after 24 h, with the decrease being more pronounced in *crr6* (Fig. 1C). The enhanced PSII photoinhibition observed in *crr6* under chilling stress is consistent with previous reports (Yamori *et al*., 2011). After recovery at 27 °C for 24 h following 24 h of chilling stress, both the active PSI amount and Fv/Fm remained lower in *crr6* than in WT (Fig. 1B–C). When Fv/Fm was plotted against the active PSI amount after 6 h of chilling stress, *crr6* exhibited a clear reduction in active PSI amount regardless of Fv/Fm, indicating PSI-specific photoinhibition (Fig. 1D). After 24 h of chilling stress, a weak positive correlation between Fv/Fm and the active PSI amount was observed, suggesting that both PSI and PSII photoinhibition (i.e. inhibition at the whole-chloroplast level) had progressed (Fig. 1E). During the recovery phase, both the active PSI amount and Fv/Fm remained lower in *crr6* than in WT (Fig. 1F). Taken together, these results demonstrate that the NDH-deficient mutant is highly susceptible to chilling-induced PSI photoinhibition, likely due to the loss of NDH-dependent Fd oxidation under chilling stress.

**Figure 1.**
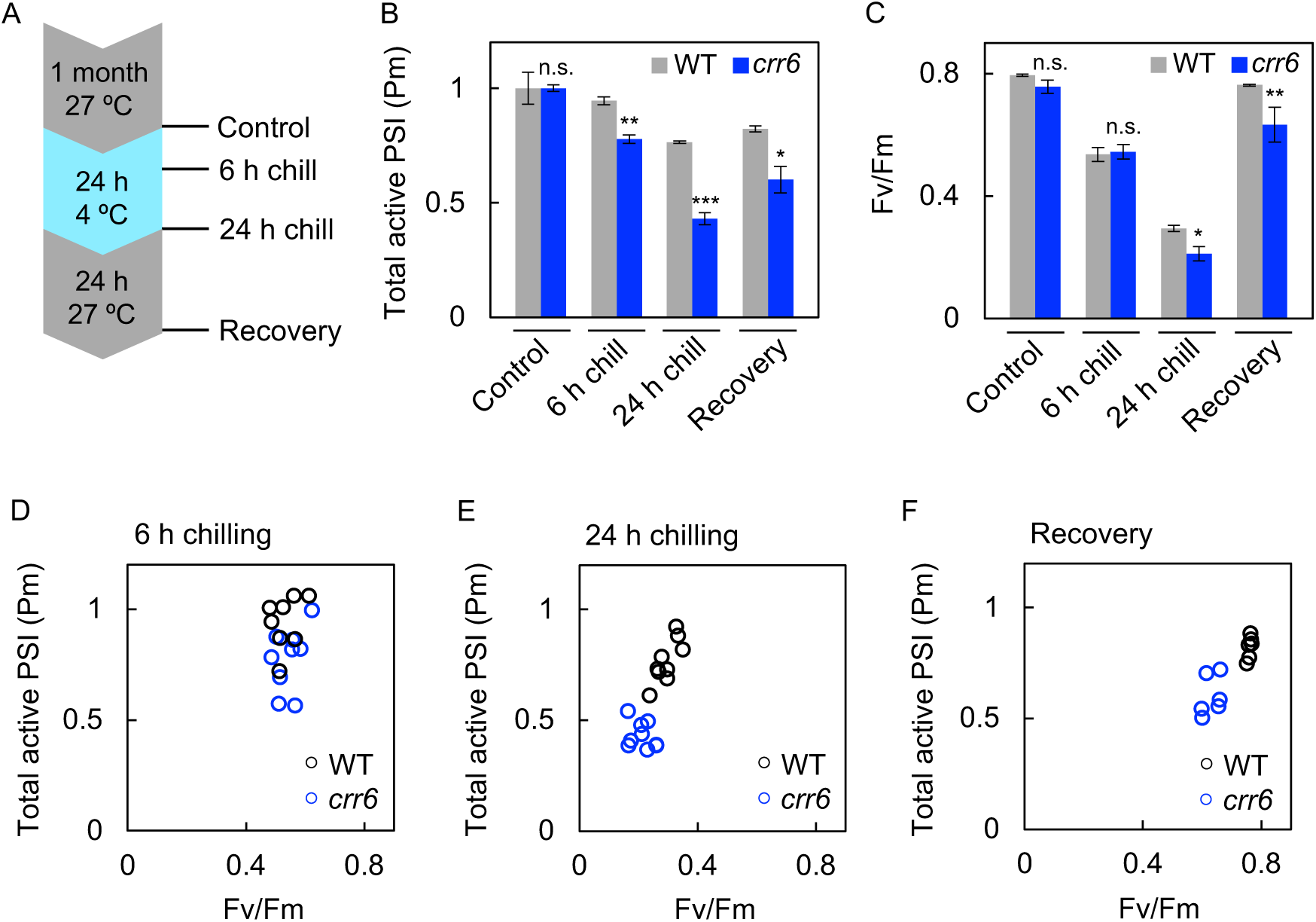
PSI and PSII photoinhibition under chilling stress in WT (Hitomebore) and NDH-deficient rice. **A**. Experimental setup for chilling stress treatment. **B–C**. Total photo-oxidizable PSI reaction center chlorophyll P700 (Pm) and the maximum quantum yield of PSII (Fv/Fm) measured before (Control) and after chilling stress. Pm values measured before chilling stress were normalized to 1. Values are the mean ± SE, n = 3, biological replicates. Asterisks (*, **, ***) indicate statistically significant differences (*p*<0.05, *p*<0.01, and *p*<0.001, respectively), whereas ‘n.s.’ denotes no significant difference (*p*>0.05) (Student’s *t*-test). **D–F**. Relationship between Fv/Fm and Pm in the same leaves analyzed in B and C.

### Electron transport activity in the NDH-deficient mutant under chilling stress

To investigate the cause of PSI photoinhibition in *crr6*, electron transport activities of PSII and PSI were examined before and after 6 h of chilling stress using Dual-PAM 100. Under control conditions before chilling stress, *crr6* showed slightly lower Y(II), a delayed increase in the oxidation level of P700 (Y(ND)), and higher Y(NA) during the induction phase of photosynthesis, indicating transient over-reduction on the PSI acceptor side, as described previously (Nikkanen *et al*., 2018; Storti *et al*., 2020; Rodriguez-Heredia *et al*., 2022; Zhou *et al*., 2023; Takeuchi *et al*., 2025*a*). These differences gradually diminished over time after the onset of light illumination, and the parameters converged to WT levels (Fig. 2A–C; Supplementary Fig. S2A). However, after 6 h of chilling stress, Y(II) and Y(ND) remained consistently lower, and Y(NA) remained higher in *crr6* than in WT (Fig. 2D–F; Supplementary Fig. S2B). Because Y(NA) correlates with the accumulation of reduced ferredoxin (Takeuchi *et al*., 2025*a*), these results indicate that loss of NDH causes electron over-accumulation at PSI.

**Figure 2.**
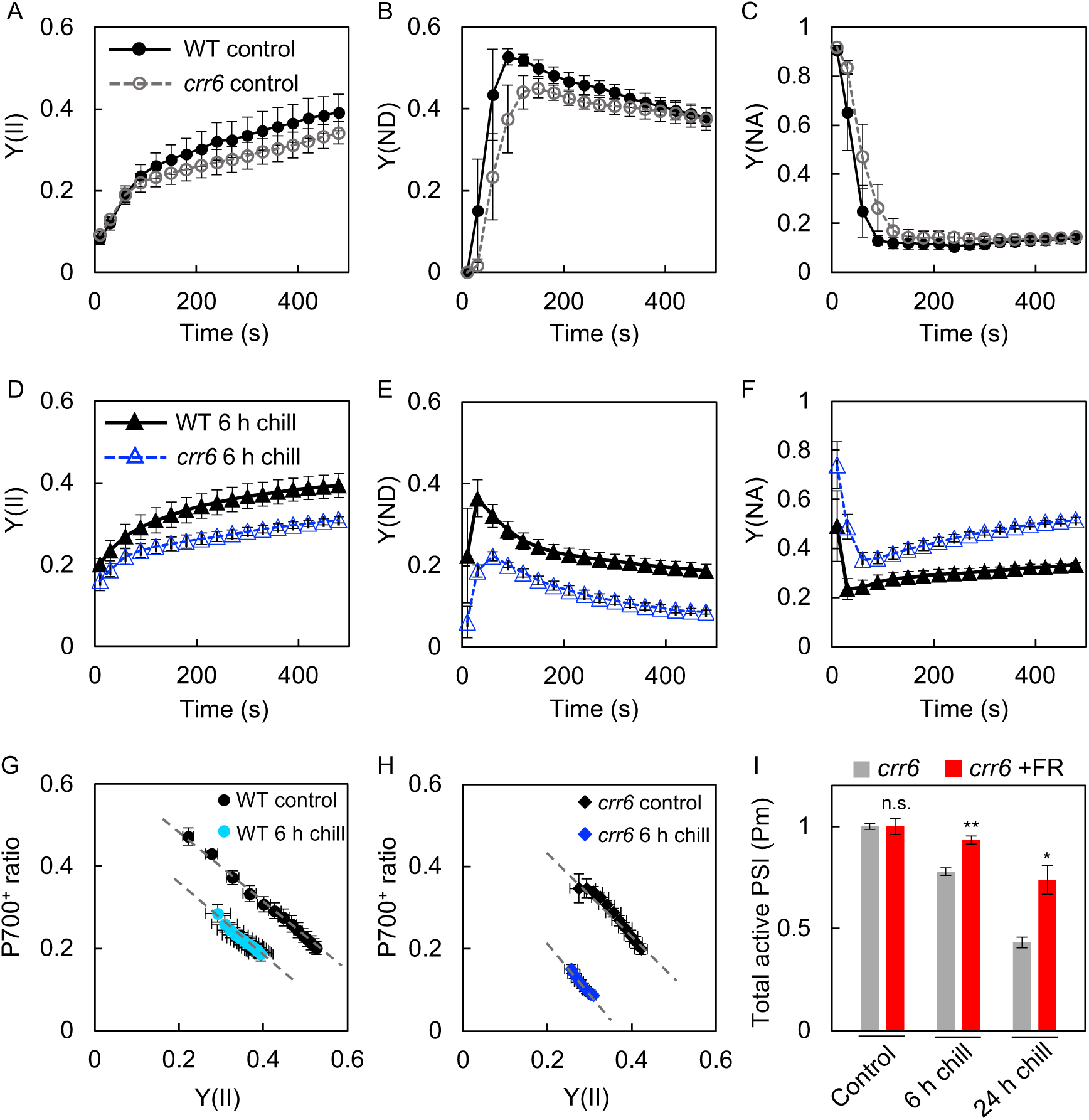
Photosynthetic activity before and after chilling stress in WT (Hitomebore) and NDH-deficient rice. **A–C**. Induction phases of Chl fluorescence and P700 redox changes analyzed using Dual-PAM 100. After at least 20 min of dark adaptation, intact leaves of WT and *crr6* were measured before chilling stress (Control). Leaves were illuminated with AL at 260 µmol photons m^−2^ s^−1^ for 8 min under ambient conditions (23 °C) to measure Y(II), the effective quantum yield of PSII; Y(ND), the ratio of P700^+^ under AL; and Y(NA), the ratio of P700 that cannot be oxidized by SP. Values are the mean ± SE, n = 3, biological replicates. **D–F**. The same measurements as shown in A–C after 6 h of chilling stress. **G–H**. Correlations between Y(II) and P700^+^ ratio (Y(ND)) before and after 6 h of chilling stress. Values were plotted using steady-state levels at 1 min after AL illumination for WT and at 2.5 min after AL illumination for *crr6*, corresponding to panels A, B, D, and E. **I**. Protective effect of FR light on PSI. Plants were subjected to light-chilling stress in the presence or absence of FR light, after which Pm was measured. Pm values of control plants were normalized to 1. Values are the mean ± SE, n = 6, biological replicates. The asterisks (*, **) indicate statistically significant differences (*p*<0.05 and *p*<0.01, respectively), whereas ‘n.s.’ denotes no significant difference (*p*>0.05) (Student’s *t*-test). FR light source was IR LED Stick (NAMOTO, Chiba, Japan) with an intensity of 50–80 µmol photons m^−2^ s^−1^.

Because P700 oxidation is inversely correlated with the rate of electron flow from PSII (Ozaki *et al*., 2023; Maekawa *et al*., 2024; Nakamura *et al*., 2024), evaluation of Y(ND) requires comparison with Y(II). Therefore, the P700⁺ ratio (Y(ND)) was plotted against the simultaneously measured Y(II). For a given Y(II), Y(ND) slightly decreased in WT after 6 h of chilling stress, it decreased much more markedly in *crr6*, indicating that PSI over-reduction occurs more readily in *crr6* (Fig. 2G–H). To further confirm that PSI over-reduction is responsible for PSI photoinhibition in *crr6*, plants were exposed to chilling stress under far-red (FR) light, which maintains PSI in an oxidized state, and the progression of photoinhibition was examined. Under FR light, PSI photoinhibition was markedly alleviated in both WT and *crr6* (Fig. 2I; Supplementary Fig. S3). This finding indicates that maintaining PSI in an oxidized state substantially suppresses chilling-stress-induced PSI photoinhibition. Collectively, the present results demonstrate that, consistent with findings in cucumber, NDH plays a crucial role in suppressing PSI over-reduction under chilling stress in rice, thereby preventing PSI photoinhibition.

### Suppression of chilling-induced PSI photoinhibition in *Flv*-expressing rice

To further test whether suppressing PSI over-reduction prevents chilling-induced PSI photoinhibition, we examined rice plants expressing *Flv*, which safely dissipate electrons downstream of PSI. Using the same experimental flow as in Fig. 1A, WT rice (cv. Jp Nipponbare) and *Flv*-expressing rice (cv. Jp Nipponbare) were grown and subjected to chilling stress for 24 h. Interestingly, in *Flv* plants, the total amount of active PSI was remained high at both 6 h and 24 h of chilling stress, indicating that PSI photoinhibition did not progress (Fig. 3A). In addition, PSII photoinhibition was also alleviated relative to WT (Fig. 3B). Plots of Fv/Fm against the total amount of active PSI further showed that, whereas PSI photoinhibition progressed in WT independently of changes in Fv/Fm, no such decrease in active PSI occurred in *Flv* plants (Fig. 3C–E). During chilling stress, *Flv* plants exhibited higher Y(II) and Y(ND) and maintained lower Y(NA) than WT plants (Fig. 3F–H; Supplementary Fig. S4–S5). Moreover, the relationship between Y(II) and the P700⁺ ratio revealed that, although the P700⁺ ratio at a given Y(II) decreased under chilling stress in WT plants, it remained largely unchanged in *Flv* plants (Fig. 3I–J), indicating that PSI over-reduction was effectively suppressed by *Flv* during chilling stress. These results demonstrate that, as observed for NDH, protection of the PSI acceptor side by *Flv* prevents PSI photoinhibition under chilling stress, highlighting the essential role of PSI acceptor-side oxidation in PSI protection.

**Figure 3.**
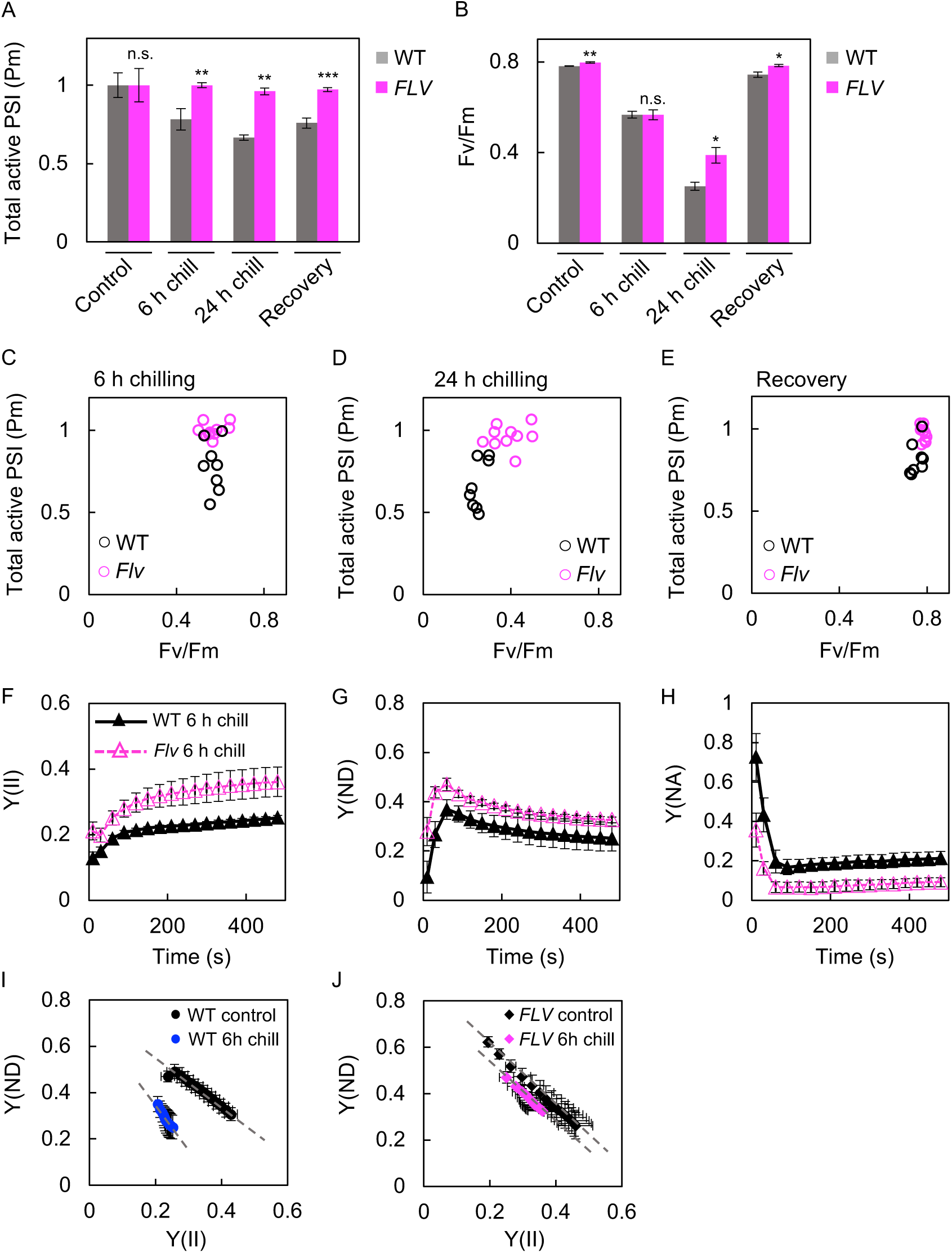
PSI and PSII photoinhibition under chilling stress in WT (Nipponbare) and *Flv*-expressing rice. **A–B**. Pm and Fv/Fm measured before (Control) and after chilling stress. Pm values of control plants were normalized to 1. Values are the mean ± SE, n = 4–5, biological replicates. The asterisks (*, **, ***) indicate statistically significant differences (*p*<0.05, *p*<0.01, and *p*<0.001, respectively), whereas ‘n.s.’ denotes no significant difference (*p*>0.05) (Student’s *t*-test). **C–E**. Relationship between Fv/Fm and Pm in the same leaves analyzed in A and B. **F–H**. Induction phases of Chl fluorescence and P700 redox changes analyzed using Dual-PAM 100. After at least 20 min of dark adaptation, intact leaves of WT and *Flv*-expressing plants were measured after 6 h of chilling stress. Leaves were illuminated with AL at 260 µmol photons m^−2^ s^−1^ for 8 min under ambient conditions (23 °C) to measure Y(II), Y(ND), and Y(NA). Values are the mean ± SE, n = 4, biological replicates. **I–J**. Correlations between Y(II) and P700^+^ ratio (Y(ND)) before and after 6 h of chilling stress. Values were plotted using steady-state levels at 1 min after AL illumination for WT and 30 s for *Flv*-expressing plants, corresponding to panels F, G, and Supplementary Fig. S4.

### Mechanism of NDH inactivation under chilling stress

The results above demonstrated the importance of NDH in chilling-induced PSI photoinhibition. We next addressed our second question, namely how NDH becomes inactivated under chilling stress, using wild-type cucumber plants, which exhibit pronounced PSI photoinhibition under chilling conditions.

To confirm chilling-induced PSI photoinhibition in cucumber WT plants, cucumbers were subjected to chilling stress for 3 h. This treatment resulted in a decrease in active PSI, followed by leaf bleaching (Fig. 4A–B). NDH activity, which is characteristically detected as a transient increase in chlorophyll (Chl) fluorescence after actinic light (AL) is turned off (Shikanai *et al*., 1998), was measured in these plants after 1 h of chilling stress, revealing complete loss of NDH activity (Fig. 4C). Moreover, even after recovery at 27 °C for 24 h following 1 h of chilling stress, NDH activity remained lower than that of control plants, suggesting that, similar to PSI, NDH activity requires prolonged recovery (Fig. 4C). In contrast, when chilling stress was applied in the dark, no decrease in NDH activity was observed (Fig. 4C), indicating that the loss of NDH activity is caused by photo-oxidative stress. In addition, electron flow from Fd to PQ during chilling stress was examined using an *in vitro* CEF assay with isolated thylakoid membranes. In thylakoids isolated before chilling treatment, inhibition of non-NDH-mediated CEF by antimycin A (AA) reduced CEF activity compared with that observed in the absence of AA (Fig. 4D) (Endo *et al*., 1998; Ueda *et al*., 2012; Courteille *et al*., 2013; Yamamoto *et al*., 2021; Nakamura *et al*., 2026). By contrast, in thylakoids isolated after 1 h of chilling stress, AA caused a much larger reduction in CEF activity (Fig. 4D), indicating that NDH-dependent CEF is strongly suppressed under chilling stress.

**Figure 4.**
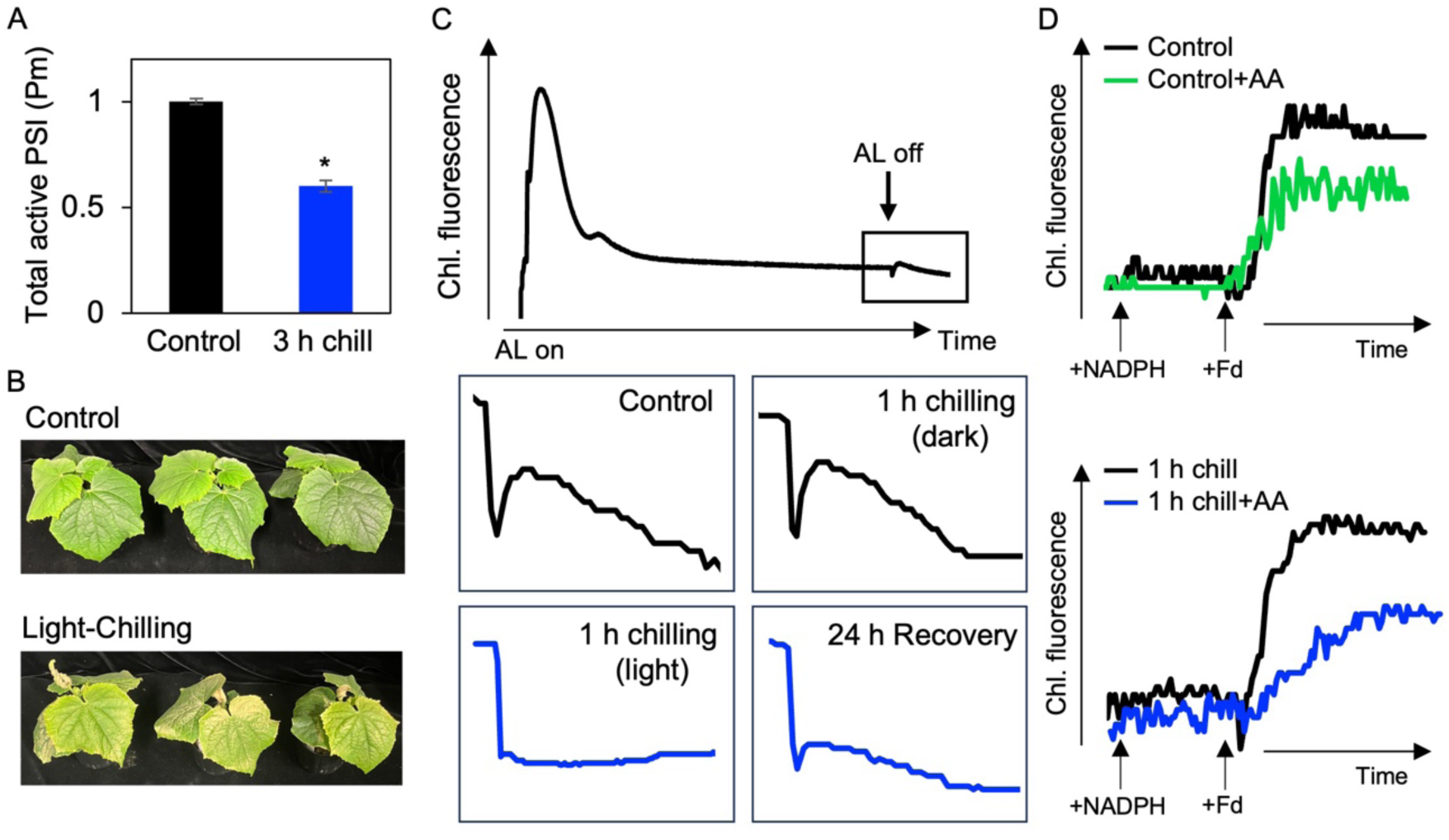
PSI photoinhibition and NDH activity under chilling stress in cucumber. **A**. Pm measured spectroscopically using Dual-PAM 100 before (Control) and after 3 h of chilling stress. **B**. Phenotype of cucumbers (cv HG) subjected to light-chilling stress for 12 h. After chilling stress, plants were returned to the original chamber at 27 °C for 2 days. **C**. Analysis of post-illumination fluorescence rise (PIFR). Cucumbers were treated at 4 °C for 1 h under either light or dark conditions. Plants after 1 h of light-chilling stress were returned to the original chamber for 1 day of recovery. Leaves before and after chilling stress were kept in the dark for at least 20 min, followed by exposure to AL at 23 °C for 5 min. The subsequent transient increase in Chl fluorescence after switching off AL was monitored in the dark (boxed area). **D**. *In vitro* Fd-dependent PQ reduction assay. Isolated thylakoids obtained before and after 1 h of light-chilling stress (20 µg Chl mL^−1^) were analyzed following the addition of 250 µM NADPH and 2.5 µM Fd. Black traces indicate measurements without antimycin A (AA), whereas colored traces indicate those with 5 µM AA.

### Structural disassembly of the PSI–NDH supercomplex during chilling stress

To identify the cause of the decline in NDH activity under chilling stress, we examined the stability of the PSI–NDH sc before and after chilling stress using BN-PAGE followed by 2D SDS-PAGE and immunoblotting. Under normal conditions, NDH is sandwiched by two PSI units (left and right PSI) to form the intact PSI–NDH sc. Accordingly, when thylakoids isolated from cucumber were subjected to BN-PAGE, the NDH subunit NdhH was detected at band 1 and at higher-molecular-mass regions (Fig. 5A, red line) (Otani *et al*., 2018; Yamamoto *et al*., 2021). However, after 1 h of chilling stress, the NdhH signal additionally appeared at lower-molecular-mass positions corresponding to band 2 (blue line) and band 3 (green line) (Fig. 5B), suggesting structural disassembly of the PSI–NDH sc (Takeuchi *et al*., 2025*a*).

**Figure 5.**
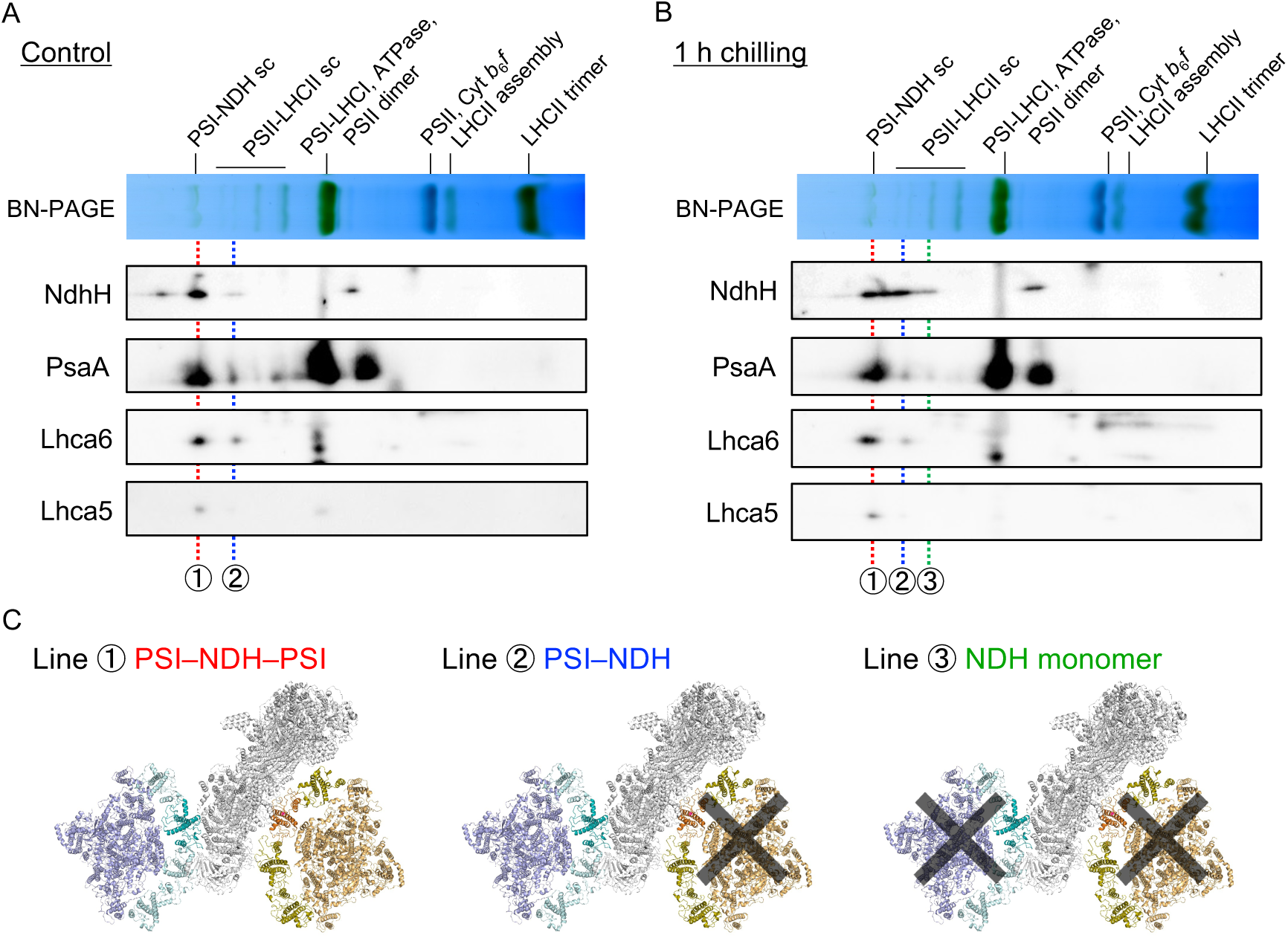
Degradation of the PSI–NDH supercomplex under chilling stress. **A–B**. Blue-native PAGE (BN-PAGE) and two-dimensional BN/SDS-PAGE analyses of thylakoid membrane proteins. Isolated thylakoids obtained before and after 1 h of light-chilling stress were solubilized with 1.0% (w/v) β-DM, and proteins corresponding to 5 µg Chl were loaded onto 4–13% gradient BN gels. Protein complexes separated by BN-PAGE were analyzed by 2D-SDS-PAGE. NDH subunit (NdhH) and PSI–LHCAsubunits (PsaA, Lhca6, and Lhca5) were detected using specific antibodies. Red dotted line 1 indicates the right PSI–NDH–left PSI sc band; blue dotted line 2 indicates the left PSI–NDH complex band; and green dotted line 3 indicates the NDH monomer. **C**. Predicted structures of the PSI–NDH sc corresponding to bands 1–3. The structures were generated based on the PSI–NDH structure (PDB ID: 7WG5).

To investigate the cause of this disassembly, immunoblot analyses were performed using antibodies against PSI core subunit PsaA, Lhca6 (which mediates the association between left PSI and NDH), and Lhca5 (which mediates the association between right PSI and NDH) (Fig. 5A–B). Band 1 (red line) contained signals for PsaA, Lhca5, and Lhca6, confirming that it represents the intact PSI–NDH sc in which NDH is bound by PSI on both sides (Fig. 5C). Band 2 (blue line) showed signals for PsaA and Lhca6 but not for Lhca5, indicating a PSI–NDH complex lacking right PSI (Fig. 5C) (Otani *et al*., 2018; Opatíková and Kouřil, 2024). The greater stability of left PSI association with NDH is consistent with previous observations that purified preparations are enriched in left-PSI–NDH complexes (Kouřil *et al*., 2014; Yadav *et al*., 2017; Su *et al*., 2022). In band 3 (green line), neither Lhca5 nor Lhca6 was detected, indicating that the Ndh signal at this position corresponds to NDH monomers dissociated from both left and right PSI under chilling stress (Fig. 5C; Supplementary Fig. S6) (Ueda *et al*., 2012). Detailed information on band positions and corresponding NDH complex states is provided in Supplementary Fig. S6 and discussed below. These results suggest that photo-oxidative stress under chilling conditions first induces dissociation of the weakly associated right PSI from NDH, followed by dissociation of left PSI mediated by Lhca6, ultimately resulting in monomerization of NDH. Because monomeric NDH is unable to function properly (Kato *et al*., 2018*a*; Otani *et al*., 2018), this disassembly likely leads to loss of NDH activity and subsequent PSI over-reduction (Fig. 4).

### Damage responsible for PSI–NDH supercomplex dissociation

Oxidative damage to Lhca has been reported during chilling-induced PSI photoinhibition (Rajagopal *et al*., 2005; Alboresi *et al*., 2009). We therefore hypothesized that PSI–NDH sc destabilization is caused by damage to Lhca proteins that support complex formation, and that degradation of Lhca6 in particular contributes to monomerization of NDH. To test this hypothesis, thylakoids were isolated from cucumber before and after chilling stress, and the total protein levels of Lhca6, Lhca3, NDH subunit PnsB1, PsaA, and Cyt *f* (control) were analyzed by immunoblotting. The abundance of Lhca6 in thylakoid membranes, a component required for PSI–Lhca–NDH sc assembly, was reduced after chilling stress (Fig. 6A–B). In contrast, no significant changes were observed in the thylakoid levels of Lhca3, PnsB1 and PsaA. To further assess damage to Lhca, we examined changes in FR light absorption mediated by Lhca (Takagi *et al*., 2026). FR light is predominantly absorbed by Lhca and excites P700, resulting in the formation of P700⁺ (Nelson and Junge, 2015). To eliminate effects of the PSI acceptor side, FR illumination was performed in the presence of methyl viologen (MV), an artificial PSI electron acceptor. After chilling stress, the rate of FR-induced P700⁺ accumulation was reduced (Fig. 6C), indicating Lhca impairment in cucumber, as previously reported (Havaux and Davaud, 1994; Sonoike, 1999). Consistent with this, absorption spectra of isolated thylakoids revealed a pronounced decrease in chlorophyll *b*–specific regions after chilling stress (approximately 650 nm and 450 nm), which are characteristic of Lhca-containing complexes (Supplementary Fig. S7) (Hui *et al*., 2000). These results suggest that, within the PSI–Lhca–NDH sc, Lhca proteins are damaged at an early stage under chilling stress, leading to monomerization of NDH and consequent PSI over-reduction.

**Figure 6.**
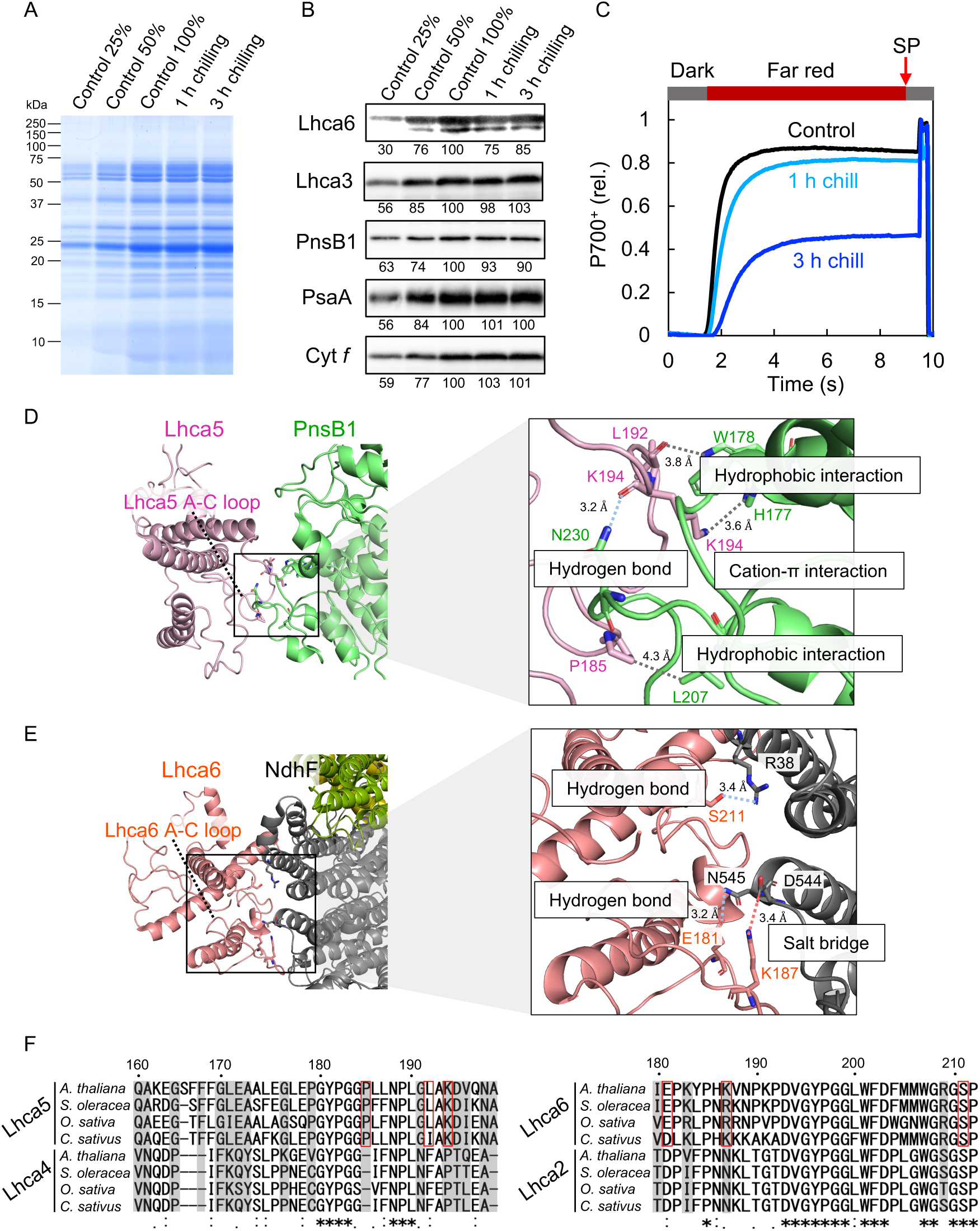
Damage to Lhca under chilling stress and interaction between PSI–LHCI and NDH. **A**. SDS-PAGE analysis of isolated thylakoid proteins before and after 1–3 h of light-chilling stress. Protein samples corresponding to 5 µg Chl (defined as 100%) were loaded onto 12% SDS-PAGE gels and stained with CBB. **B**. Immunoblot analysis of proteins separated by SDS-PAGE using specific antibodies. Numbers indicate relative band intensities quantified using ImageJ. **C**. P700 oxidation kinetics under FR light and SP before and after 1–3 h of light-chilling stress. Leaves were vacuum-infiltrated with 50 µM methyl viologen and sufficiently dark-adapted, followed by illumination 10 s FR illumination and application of a 200 ms SP using Dual-PAM 100. P700 oxidation kinetics were normalized to total photo-oxidizable P700. **D–E**. Structural views of interactions between Lhca5 and PnsB1 (D) and between Lhca6 and NdhF (E). Right panels show magnified views of the stromal A–C loop regions and predicted interaction sites. **F**. Multiple sequence alignments of Lhca5 and Lhca4, and of Lhca6 and Lhca2, generated using ClustalW. Lhca5- and Lhca6-specific residues are shaded in black, and residues predicted to be important for Lhca5–NDH interaction are outlined in red. Asterisks (*), colons (:), and dots (.) below the aligned sequences indicate identical, conserved, and semi-conserved amino acids, respectively.

### Interactions between Lhca and NDH

To clarify the cause of NDH monomerization in cucumber under chilling stress, we examined the interaction sites between Lhca and NDH and compared them across plant species. Lhca5 and Lhca6 replace Lhca4 and Lhca2, respectively, within the PSI–Lhca complex to enable PSI–NDH association, and therefore share sequence similarity with the corresponding Lhca proteins. Based on amino acid sequence and structural comparisons between Lhca5 and Lhca4, and between Lhca6 and Lhca2, we identified residues potentially important for PSI–Lhca–NDH interaction. Using previously reported cryo-EM structures of the PSI–NDH sc, we then examined the structural modes of interaction with NDH (see Supplementary Fig. S8–11 for details).

For the interaction between right PSI and NDH, contacts were identified between the stromal A–C loop of Lhca5 and PnsB1 (Fig. 6D; Supplementary Fig. S8) (Shen *et al*., 2022; Su *et al*., 2022). In this interface, hydrophobic interactions involving P185 and L192, hydrogen bonding via the main chain of K194, and a cation–π interaction via the side chain of K194 were identified as key interaction sites (Fig. 6D). For the interaction between left PSI and NDH, interactions were observed between the stromal A–C loop region of Lhca6 and NdhF (Fig. 6E; Supplementary Fig. S9) (Otani *et al*., 2017; Shen *et al*., 2022). In this interface, a salt bridge formed by the side chain of K(R)187, together with hydrogen bonds involving the side chains of E181 and S211, was identified as a key interaction site (Fig. 6E). The presence of salt bridge provides a plausible explanation for the stronger association between Lhca6 and NDH relative to that between Lhca5 and NDH (Yadav *et al*., 2017). Interestingly, analysis of residue E181 in Lhca6 revealed that, whereas this position is conserved as glutamate (E181) in most plant species, it is substituted with aspartate (D181) in cucumber (Fig. 6F; Supplementary Fig. S10–S11). This suggests that the binding strength between left PSI and NDH in cucumber may differ from that in other species. Together, these findings provide a structural basis for preferential dissociation of right PSI from NDH and the propensity for NDH monomerization in cucumber under chilling stress.

## Discussion

NDH accumulation is known to increase under chilling stress (Bernhard Teicher *et al*., 2000), yet the physiological significance of this response has remained unclear. In this study, using an NDH-deficient mutant, we demonstrate that NDH suppresses PSI over-reduction under chilling stress, thereby mitigating PSI photoinhibition. Furthermore, based on our analyses in cucumber, we propose a mechanistic framework underlying chilling-induced PSI photoinhibition (Fig. 7). (1): Suppression of the CBB cycle under chilling stress causes transient over-reduction of PSI, leading to oxidative damages to Lhca that stabilize the PSI–NDH sc (Fig. 6). (2): Damage to Lhca promotes dissociation of NDH from the PSI–Lhca, resulting in incomplete PSI–NDH assemblies and subsequent monomerization of NDH (Fig. 5; Supplementary Fig. S6). (3): Impaired electron efflux from Fd promotes PSI over-reduction, ultimately leading to PSI photoinhibition (Fig. 4, 7).

**Figure 7.**
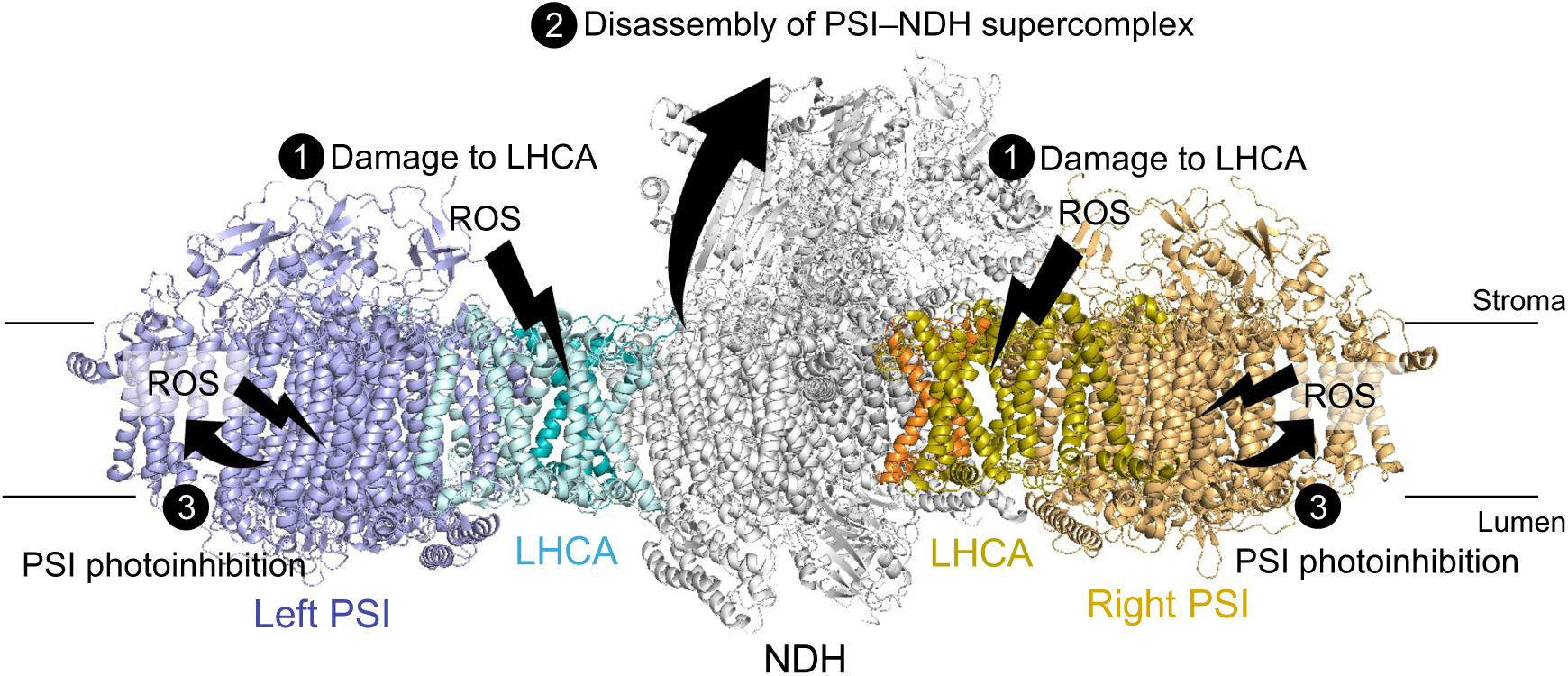
Model for PSI photoinhibition under chilling stress. Under chilling stress, reduced activity of the CBB cycle causes over-reduction of PSI, resulting in ROS production at PSI and Lhca. (1) ROS induces oxidative damage to Lhca, which is essential for maintaining the PSI–NDH sc, thereby destabilizing the complex. (2) The two PSI–LHCI units dissociate from the NDH complex, leading to monomerization of NDH and loss of NDH activity. (3) Reduced NDH activity further enhances over-reduction of PSI, promotes ROS generation near the PSI core, and ultimately leads to PSI photoinhibition.

Lhca has been identified as a target of PSI photoinhibition (Lhca photoinhibition) (Takagi *et al*., 2026). When excessive light limits excitation energy transfer from Lhca to the PSI core, triplet chlorophyll (^3^Chl) is formed within Lhca, leading to the generation of ^1^O_2_ (Carbonera *et al*., 2005; Rajagopal *et al*., 2005; Kale *et al*., 2020). Consistent with this, analyses of Lhca from field-grown plants have revealed ROS-oxidized amino acid residues located near chlorophyll molecules and on the protein surface (Kale *et al*., 2020, 2022). These observations suggest that PSI over-reduction under chilling stress may impair excitation energy transfer within Lhca, thereby promoting ^1^O_2_ production within Lhca and leading to its degradation. Although Lhca possesses carotenoid-dependent excitation energy quenching mechanisms, these protective mechanisms become insufficient under excessive stress, leading to oxidative damage (Croce *et al*., 2007; Alboresi *et al*., 2009; Cazzaniga *et al*., 2012, 2016; Ballottari *et al*., 2014; Azai *et al*., 2020). Furthermore, because the interactions between Lhca5 or Lhca6 and NDH are mediated by stromal loop regions (Fig. 6), these interfaces are particularly susceptible to ROS attack (Kale *et al*., 2020). Such ROS-induced oxidative damage to Lhca likely destabilizes the Lhca–NDH interaction. Formation of the PSI–NDH sc via Lhca has been proposed to protect NDH from photodamage (Kato *et al*., 2018*a*); therefore, destabilization of the PSI–NDH sc caused by Lhca damage at the early phase of chilling stress is likely a major cause of NDH inactivation and consequent PSI over-reduction.

The physiological roles of NDH have been progressively elucidated. In C_3_ plants, NDH has been shown *in vivo* to function in ferredoxin oxidation, reduction of PQ pool, and electron sink activity within its internal Fe–S clusters, while *in vitro* studies and analyses in cyanobacteria have demonstrated its proton-pumping capability (Strand *et al*., 2017; Miller *et al*., 2021; Richardson *et al*., 2021; Zhang *et al*., 2025). Regulation of NDH activity by oxidative stress and the thioredoxin system supports its role in suppressing ROS formation at PSI (Lascano *et al*., 2003; Wang *et al*., 2006; Nikkanen *et al*., 2018; Pan *et al*., 2020; Ermakova *et al*., 2024; Zheng *et al*., 2025). In addition, NDH has been implicated in the regulation of plastid-associated metabolic processes (Nashilevitz *et al*., 2010; Zhang *et al*., 2023, Zhang Q *et al*., 2024), most likely through alterations in NADPH metabolism via NDH (Kambakam *et al*., 2016; Nikkanen *et al*., 2020). In the present study, we demonstrate that NDH plays a crucial role in suppressing PSI over-reduction under chilling stress (Fig. 1–2). This finding is consistent with recent reports that NDH functions as an electron sink and alleviates PSI photoinhibition when the CBB cycle is restricted (Krämer *et al*., 2024; Takeuchi *et al*., 2025*a*; Wang *et al*., 2026). In addition, NDH-mediated reduction of the PQ pool can suppress electron transfer at the cytochrome *b*_6_*f* complex independently of ΔpH, thereby restricting electron flow toward PSI and preventing PSI over-reduction under chilling stress (Shaku et al., 2016; Tikhonov, 2024). Furthermore, our results reveal that maintenance of the PSI–NDH sc is critical for sustaining NDH activity under stress conditions (Fig. 5; Fig. 7; Supplementary Fig. S6). Structural alterations of the PSI–NDH sc are therefore likely under various stress conditions in which changes in NDH activity have been reported (Martín *et al*., 2004; Sun *et al*., 2017; Wu *et al*., 2019; Wang *et al*., 2020; Zhang *et al*., 2023; He *et al*., 2024). NDH is prone to light-induced degradation in the absence of Lhca6, suggesting that Lhca6 was evolutionarily acquired to stabilize NDH under illuminated conditions (Kato *et al*., 2018*a*). Our findings further support the evolutionary significance of the acquisition of Lhca4 and Lhca6 during the transition from bryophytes to vascular plants, which enabled stabilization of the PSI–NDH sc and maintenance of NDH function under stress (Kato *et al*., 2018*a*; Opatíková and Kouřil, 2024).

Multiple forms of the PSI–NDH sc were identified in immunoblot analyses following 2D BN/SDS-PAGE (Fig. 5; Supplementary Fig. S6). NDH detected in higher-molecular-mass regions than band 1 (red line) exclusively under control conditions corresponds to a PSI–NDH sc in which more than two PSI units are associated with left PSI (Fig. 5A) (Yadav *et al*., 2017). In addition, band 2 (blue line), representing a PSI–NDH complex lacking right PSI, was detected even under control conditions (Fig. 5A; Supplementary Fig. S6), supporting the intrinsic instability of right PSI (Kouřil *et al*., 2014; Yadav *et al*., 2017). Interestingly, the right-PSI binding site may accommodate NDH regulators or interactions with the cytochrome *b*_6_*f* complex (Li *et al*., 2018; Su *et al*., 2022; He *et al*., 2026), raising the possibility that differential binding capacities of left and right PSI to NDH confer functional significance beyond structural stability. Recently, we reported that PIFI contributes to the stability of the PSI–NDH supercomplex; however, clarification of the binding site of PIFI on NDH and the detailed mechanism by which it regulates NDH activity will be required (Kohzuma *et al*., 2026).

Multiple assembly intermediates of NDH have been reported. We revealed NdhH signals in a lower-molecular-mass region than those corresponding to the NDH monomer and the PSI–LHCI complex (Supplementary Fig. S6, magenta line). At this position, signals were also detected using antibodies against NdhT (a component of SubE) and PnsL (a component of SubL), in addition to NdhH (a component of SubA). In contrast, no signal was detected at this position with antibodies against PnsB1, a component of SubB (Supplementary Fig. S6, magenta line). These observations indicate that the low-molecular-mass signal represents an assembly intermediate composed of SubA, SubE, and SubL (Armbruster *et al*., 2013; Otani *et al*., 2018; Ishikawa *et al*., 2020). PsaA signal was also detected at this position (magenta line), whereas no Lhca signal was observed, suggesting that this band corresponds to a PSI core monomer lacking Lhca association (Supplementary Fig. S6). In addition, immunoblotting with anti-PsaA antibodies revealed a faint signal near band 3 (green line), corresponding to the NDH monomer position (around 770 kDa) (Fig. 5A; Supplementary Fig. S6). This position closely matches the reported molecular mass of a SubB–PSI–Lhca assembly intermediate (around 750 kDa) (Armbruster *et al*., 2013). Consistent with this, a signal at band 3 (green line) was detected only with antibodies against PnsB1 (a component of SubB) (Supplementary Fig. S6), indicating that this band represents a SubB–PSI assembly intermediate (Kato *et al*., 2018*b*; Ishikawa *et al*., 2020). Although mechanistic analyses were conducted in cucumber, the conservation of PSI–NDH architecture among angiosperms suggests that similar structural vulnerability may operate in rice.

In this study, analyses of chilling responses in rice revealed three key findings. 1: Even WT rice undergoes PSI photoinhibition under chilling stress (Fig. 1–3), and loss of NDH results in markedly more severe PSI photoinhibition (Fig. 1). 2: FR light effectively suppresses chilling-induced PSI photoinhibition (Fig. 2I). 3: FLV confers strong protection against chilling-induced PSI photoinhibition (Fig. 3). It has been reported that growth of NDH-deficient mutants is impaired under chilling conditions (Yamori *et al*., 2011; Urzinger *et al*., 2025). Our results demonstrate that PSI photoinhibition is a major cause of this sensitivity. It has also been reported that introduction of FLV into angiosperms enhances tolerance to fluctuating light, salt stress, and drought stress (Yamamoto *et al*., 2016; Wada *et al*., 2018; Vicino *et al*., 2021; Suganami *et al*., 2022). Our findings extend these observations by demonstrating that FLV also effectively protect PSI under chilling stress. Notably, needles of conifers such as Scots pine and Norway spruce, which have lost NDH during evolution (Wakasugi *et al*., 1994; Braukmann *et al*., 2009), rely on FLV to alleviate early-spring chilling damage (Bag *et al*., 2023). This suggests functional complementarity between NDH and FLV in PSI protection. Thus, in field-grown angiosperms lacking FLV, suppression of PSI over-reduction by the PSI–NDH sc, together with enhancement of P700 oxidation by FR light (Kono *et al*., 2017, 2022), abundant in natural sunlight, provides effective protection against chilling-induced PSI photodamage in natural environments.

## Conclusion

This study demonstrates that NDH contributes to chilling stress tolerance by suppressing PSI photoinhibition. In particular, maintaining the PSI–NDH sc under chilling conditions is essential for NDH function, and damage to Lhca leads to NDH inactivation by destabilizing the PSI–NDH sc. Because both the abundance of NDH and the architecture of the PSI–NDH sc vary among plant species and tissues (Laihonen et al., 2024), our findings in cucumber highlight the importance of extending NDH research beyond model species. Further studies using diverse plant species will likely uncover additional, previously unrecognized roles of NDH in photosynthetic stress tolerance.

## Acknowledgments

We sincerely thank Dr. Masaru Kono (Astrobiology Center, Japan) for valuable discussions and insightful suggestions, and Dr. Amane Makino and Dr. Shinya Wada for kindly providing seeds of the *crr6* and *FLV* lines. This work was supported in part by CREST, Japan Science and Technology Agency (JPMJCR15O3 and JPMJCR17O2 to K.I.) and by Grant-in-Aid for JSPS Fellows (JP23KJ1357 to K.T.).

## Author contributions

KI conceived the project. KT, SH, and KI designed the research. KT and SH performed the experiments. KT and SH drafted the original manuscript. KT and KI revised and finalized the manuscript.

## Data availability

The data underlying this article are available in the article.

## Supporting Information

**Supplementary Fig. S1.** Post-illumination fluorescence rise (PIFR) analyzed under ambient conditions (23 °C) in WT (Hitomebore) and *crr6*.

**Supplementary Fig. S2.** Photosynthetic activity before and after chilling stress in WT (Hitomebore) and *crr6*.

**Supplementary Fig. S3.** Protective effects of FR light on PSI and PSII.

**Supplementary Fig. S4.** Photosynthetic activity before chilling stress (Control) in WT (Nipponbare) and *Flv*-expressing rice.

**Supplementary Fig. S5.** Photosynthetic activity after 6 h of chilling stress in WT (Nipponbare) and *Flv*-expressing rice.

**Supplementary Fig. S6.** Analysis of the PSI–NDH sc assembly state in cucumber.

**Supplementary Fig. S7.** Absorption spectra of isolated thylakoids before and after chilling stress in cucumber.

**Supplementary Fig. S8**. Interaction sites between Lhca5 and PnsB1.

**Supplementary Fig. S9.** Interaction sites between Lhca6 and NdhF.

**Supplementary Fig. S10.** Comparison of amino acid sequences in the stromal loop regions of Lhca4 and Lhca5.

**Supplementary Fig. S11.** Comparison of amino acid sequences in the stromal loop regions of Lhca2 and Lhca6.

## References

Alboresi A, Ballottari M, Hienerwadel R, Giacometti GM, Morosinotto T. 2009. Antenna complexes protect photosystem I from photoinhibition. BMC Plant Biology 9, 71.

Allen DJ, Ort DR. 2001. Impacts of chilling temperatures on photosynthesis in warm-climate plants. Trends in Plant Science 6, 36–42.

Amin B, Atif MJ, Kandegama W, Nasar J, Alam P, Fang Z, Cheng Z. 2024. Low temperature and high humidity affect dynamics of chlorophyll biosynthesis and secondary metabolites in Cucumber. BMC plant biology 24, 903.

Armbruster U, Rühle T, Kreller R, Strotbek C, Zühlke J, Tadini L, Blunder T, Hertle AP, Qi Y, Rengstl B. 2013. The photosynthesis affected mutant68–like protein evolved from a PSII assembly factor to mediate assembly of the chloroplast NAD(P)H dehydrogenase complex in *Arabidopsis*. The Plant Cell 25, 3926–3943.

Azai C, Harada J, Fujimoto S, Masuda S, Kosumi D. 2020. Anaerobic energy dissipation by glycosylated carotenoids in the green sulfur bacterium *Chlorobaculum tepidum*. Journal of Photochemistry and Photobiology A: Chemistry 403, 112828.

Bag P, Shutova T, Shevela D, Lihavainen J, Nanda S, Ivanov AG, Messinger J, Jansson S. 2023. Flavodiiron-mediated O_2_ photoreduction at photosystem I acceptor-side provides photoprotection to conifer thylakoids in early spring. Nature Communications 14, 3210.

Ballottari M, Alcocer MJ, D’Andrea C, Viola D, Ahn TK, Petrozza A, Polli D, Fleming GR, Cerullo G, Bassi R. 2014. Regulation of photosystem I light harvesting by zeaxanthin. Proceedings of the National Academy of Sciences 111, 2431–2438.

Bernhard Teicher H, Lindberg Møller B, Vibe Scheller H. 2000. Photoinhibition of photosystem I in field-grown barley (*Hordeum vulgare* L.): induction, recovery and acclimation. Photosynthesis Research 64, 53–61.

Braukmann TWA, Kuzmina M, Stefanović S. 2009. Loss of all plastid ndh genes in Gnetales and conifers: extent and evolutionary significance for the seed plant phylogeny. Current Genetics 55, 323–337.

Carbonera D, Agostini G, Morosinotto T, Bassi R. 2005. Quenching of chlorophyll triplet states by carotenoids in reconstituted Lhca4 subunit of peripheral light-harvesting complex of photosystem I. Biochemistry 44, 8337–8346.

Cazzaniga S, Bressan M, Carbonera D, Agostini A, Dall’Osto L. 2016. Differential roles of carotenes and xanthophylls in photosystem I photoprotection. Biochemistry 55, 3636–3649.

Cazzaniga S, Li Z, Niyogi KK, Bassi R, Dall’Osto L. 2012. The Arabidopsis *szl1* mutant reveals a critical role of β-carotene in photosystem I photoprotection. Plant Physiology 159, 1745–1758.

Che Y, Kusama S, Matsui S, Suorsa M, Nakano T, Aro E-M, Ifuku K. 2020. Arabidopsis PsbP-like protein 1 facilitates the assembly of the photosystem II supercomplexes and optimizes plant fitness under fluctuating light. Plant and Cell Physiology 61, 1168–1180.

Chen S, Zheng Q, Qi Z, Ding J, Song X, Xia X. 2024. Stress-induced delay of the IP rise of the fast chlorophyll a fluorescence transient in tomato. Scientia Horticulturae 326, 112741.

Cherepanov DA, Milanovsky GE, Petrova AA, Tikhonov AN, Semenov AY. 2017. Electron transfer through the acceptor side of photosystem I: Interaction with exogenous acceptors and molecular oxygen. Biochemistry (Moscow) 82, 1249–1268.

Courteille A, Vesa S, Sanz-Barrio R, Cazalé A-C, Becuwe-Linka N, Farran I, Havaux M, Rey P, Rumeau D. 2013. Thioredoxin m4 controls photosynthetic alternative electron pathways in *Arabidopsis*. Plant Physiology 161, 508–520.

Croce R, Mozzo M, Morosinotto T, Romeo A, Hienerwadel R, Bassi R. 2007. Singlet and triplet state transitions of carotenoids in the antenna complexes of higher-plant photosystem I. Biochemistry 46, 3846–3855.

Degen GE, Johnson MP. 2024. Photosynthetic control at the cytochrome *b*_6_*f* complex. The Plant Cell 36, 4065–4079.

Endo T, Shikanai T, Sato F, Asada K. 1998. NAD(P)H dehydrogenase-dependent, antimycin A-sensitive electron donation to plastoquinone in tobacco chloroplasts. Plant and Cell Physiology 39, 1226–1231.

Erling Tjus S, Lindberg Møller B, Scheller HV. 1999. Photoinhibition of photosystem I damages both reaction centre proteins PSI-A and PSI-B and acceptor-side located small photosystem I polypeptides. Photosynthesis Research 60, 75–86.

Erling Tjus S, Lindberg Møller B, Vibe Scheller H. 1998. Photosystem I is an early target of photoinhibition in barley illuminated at chilling temperatures. Plant Physiology 116, 755–764.

Ermakova M, Woodford R, Fitzpatrick D, Nix SJ, Zwahlen SM, Farquhar GD, von Caemmerer S, Furbank RT. 2024. Chloroplast NADH dehydrogenase-like complex-mediated cyclic electron flow is the main electron transport route in C_4_ bundle sheath cells. New Phytologist 243, 2187–2200.

Foyer C, Furbank R, Harbinson J, Horton P. 1990. The mechanisms contributing to photosynthetic control of electron transport by carbon assimilation in leaves. Photosynthesis Research 25, 83–100.

Furutani R, Ohnishi M, Mori Y, Wada S, Miyake C. 2022. The difficulty of estimating the electron transport rate at photosystem I. Journal of Plant Research 135, 565–577.

Govindachary S, Bukhov NG, Joly D, Carpentier R. 2004. Photosystem II inhibition by moderate light under low temperature in intact leaves of chilling-sensitive and-tolerant plants. Physiologia Plantarum 121, 322–333.

Grebe S, Porcar-Castell A, Riikonen A, Paakkarinen V, Aro E-M. 2024. Accounting for photosystem I photoinhibition sheds new light on seasonal acclimation strategies of boreal conifers. Journal of Experimental Botany 75, 3973–3992.

Hanawa H, Ishizaki K, Nohira K, Takagi D, Shimakawa G, Sejima T, Shaku K, Makino A, Miyake C. 2017. Land plants drive photorespiration as higher electron-sink: Comparative study of post-illumination transient O_2_-uptake rates from liverworts to angiosperms through ferns and gymnosperms. Physiologia Plantarum 161, 138–149.

Hani U, Naranjo B, Shimakawa G, Espinasse C, Vanacker H, Sétif P, Rintamäki E, Issakidis-Bourguet E, Krieger-Liszkay A. 2024. A complex and dynamic redox network regulates oxygen reduction at photosystem I in *Arabidopsis*. Plant Physiology 197, kiae501.

Havaux M, Davaud A. 1994. Photoinhibition of photosynthesis in chilled potato leaves is not correlated with a loss of photosystem-II activity: preferential inactivation of photosystem I. Photosynthesis Research 40, 75–92.

He Y, Lu C, Jiang Z, Sun Y, Liu H, Yin Z. 2024. NADH dehydrogenase-like complex L subunit improves salt tolerance by enhancing photosynthetic electron transport. Plant Physiology and Biochemistry 207, 108420.

He Y, Li R, Li J, Zheng S, Jiang D, Li X, Liu Q, Zhou X, Lu J, Chen J. 2026. OsRNase Z^S2^ degrades the transcripts of NdhD and NdhF subunits of the NDH complex to modulate the balance between cyclic and linear photosynthetic electron transport at different developmental stages. Plant Communications. 10.1016/j.xplc.2026.101791.

Helman Y, Tchernov D, Reinhold L, Shibata M, Ogawa T, Schwarz R, Ohad I, Kaplan A. 2003. Genes encoding A-type flavoproteins are essential for photoreduction of O_2_ in cyanobacteria. Current Biology 13, 230–235.

Huang W, Yang Y-J, Zhang S-B. 2019. The role of water-water cycle in regulating the redox state of photosystem I under fluctuating light. Biochimica et Biophysica Acta (BBA)-Bioenergetics 1860, 383–390.

Hui Y, Jie W, Carpentier R. 2000. Degradation of the photosystem I complex during photoinhibition. Photochemistry and Photobiology 72, 508–512.

Ifuku K, Endo T, Shikanai T, Aro E-M. 2011. Structure of the chloroplast NADH dehydrogenase-like complex: nomenclature for nuclear-encoded subunits. Plant and Cell Physiology 52, 1560–1568.

Introini B, Hahn A, Kühlbrandt W. 2025. Cryo-EM structure of the NDH–PSI–LHCI supercomplex from *Spinacia oleracea*. Nature Structural & Molecular Biology 32, 968–978.

Ishikawa N, Yokoe Y, Nishimura T, Nakano T, Ifuku K. 2020. PsbQ-like protein 3 functions as an assembly factor for the chloroplast NADH dehydrogenase-like complex in *Arabidopsis*. Plant and Cell Physiology 61, 1252–1261.

Ivanov AG, Morgan RM, Gray GR, Velitchkova MY, Huner NPA. 1998. Temperature/light dependent development of selective resistance to photoinhibition of photosystem I. FEBS Letters 430, 288–292.

Järvi S, Suorsa M, Paakkarinen V, Aro E-M. 2011. Optimized native gel systems for separation of thylakoid protein complexes: novel super-and mega-complexes. Biochemical Journal 439, 207–214.

Kale R, Sallans L, Frankel LK, Bricker TM. 2020. Natively oxidized amino acid residues in the spinach PS I-LHC I supercomplex. Photosynthesis Research 143, 263–273.

Kale RS, Seep JL, Sallans L, Frankel LK, Bricker TM. 2022. Oxidative modification of LHC II associated with photosystem II and PS I-LHC I-LHC II membranes. Photosynthesis research 152, 261–274.

Kambakam S, Bhattacharjee U, Petrich J, Rodermel S. 2016. PTOX mediates novel pathways of electron transport in etioplasts of *Arabidopsis*. Molecular plant 9, 1240–1259.

Kato Y, Odahara M, Fukao Y, Shikanai T. 2018*a*. Stepwise evolution of supercomplex formation with photosystem I is required for stabilization of chloroplast NADH dehydrogenase-like complex: Lhca5-dependent supercomplex formation in *Physcomitrella patens*. The Plant Journal 96, 937–948.

Kato Y, Odahara M, Shikanai T. 2021. Evolution of an assembly factor-based subunit contributed to a novel NDH-PSI supercomplex formation in chloroplasts. Nature communications 12, 3685.

Kato Y, Sugimoto K, Shikanai T. 2018*b*. NDH-PSI supercomplex assembly precedes full assembly of the NDH complex in chloroplast. Plant Physiology 176, 1728–1738.

Kim J, Kim S, Cho SH, Chow WS, Lee C. 2005. Photosystem I acceptor side limitation is a prerequisite for the reversible decrease in the maximum extent of P700 oxidation after short-term chilling in the light in four plant species with different chilling sensitivities. Physiologia Plantarum 123, 100–107.

Kim S-J, Lee C-H, Hope A, Chow WS. 2001. Inhibition of photosystems I and II and enhanced back flow of photosystem I electrons in cucumber leaf discs chilled in the light. Plant and Cell Physiology 42, 842–848.

Kingston-Smith AH, Harbinson J, Williams J, Foyer CH. 1997. Effect of chilling on carbon assimilation, enzyme activation, and photosynthetic electron transport in the absence of photoinhibition in maize leaves. Plant Physiology 114, 1039–1046.

Klughammer C, Schreiber U. 1994. An improved method, using saturating light pulses, for the determination of photosystem I quantum yield via P700^+^-absorbance changes at 830 nm. Planta 192, 261–268.

Klughammer C, Schreiber U. 2008. Saturation Pulse method for assessment of energy conversion in PS I. PAM Application Notes 1, 11–14.

Kohzuma K, Murai M, Imaizumi K, Miura K, Kimura A, Yoshida K, Che Y, Ishikawa N, Hisabori T, Ifuku K. 2026. PIFI stabilizes chloroplast NDH-PSI supercomplex to maintain plastoquinone redox balance and PSII efficiency. bioRxiv. 10.64898/2026.03.22.713156.

Kono M, Oguchi R, Terashima I. 2022. Photoinhibition of PSI and PSII in nature and in the laboratory: Ecological approaches. Progress in Botany 84, 241–292.

Kono M, Yamori W, Suzuki Y, Terashima I. 2017. Photoprotection of PSI by far-red light against the fluctuating light-induced photoinhibition in *Arabidopsis thaliana* and field-grown plants. Plant and Cell Physiology 58, 35–45.

Kornyeyev D, Logan BA, Allen RD, Holaday AS. 2003. Effect of chloroplastic overproduction of ascorbate peroxidase on photosynthesis and photoprotection in cotton leaves subjected to low temperature photoinhibition. Plant Science 165, 1033–1041.

Kouřil R, Strouhal O, Nosek L, Lenobel R, Chamrád I, Boekema EJ, Šebela M, Ilík P. 2014. Structural characterization of a plant photosystem I and NAD(P)H dehydrogenase supercomplex. The Plant Journal 77, 568–576.

Kozuleva MA, Ivanov BN. 2010. Evaluation of the participation of ferredoxin in oxygen reduction in the photosynthetic electron transport chain of isolated pea thylakoids. Photosynthesis Research 105, 51–61.

Kozuleva M, Petrova A, Milrad Y, Semenov A, Ivanov B, Redding KE, Yacoby I. 2021. Phylloquinone is the principal Mehler reaction site within photosystem I in high light. Plant Physiology 186, 1848–1858.

Krämer M, Blanco NE, Penzler J-F, Davis GA, Brandt B, Leister D, Kunz H-H. 2024. Cyclic electron flow compensates loss of PGDH3 and concomitant stromal NADH reduction. Scientific Reports 14, 29274.

Kudoh H, Sonoike K. 2002. Irreversible damage to photosystem I by chilling in the light: cause of the degradation of chlorophyll after returning to normal growth temperature. Planta 215, 541–548.

Laihonen L, Rantala M, Ranasinghe U, Tyystjärvi E, Mulo P. 2024. Transcriptomic and proteomic analyses of distinct Arabidopsis organs reveal high PSI-NDH complex accumulation in stems. Physiologia Plantarum 176, e14227.

Lascano HR, Casano LM, Martın M, Sabater B. 2003. The activity of the chloroplastic Ndh complex is regulated by phosphorylation of the NDH-F subunit. Plant Physiology 132, 256–262.

Li L, Aro E-M, Millar AH. 2018. Mechanisms of photodamage and protein turnover in photoinhibition. Trends in Plant Science 23, 667–676.

Li J, Zhang H, Yue D, Chen S, Yin Y, Zheng C, Chen Y. 2024. Endogenous serotonin induced by cold acclimation increases cold tolerance by reshaping the MEL/ROS/RNS redox network in *Kandelia obovata*. Journal of Forestry Research 35, 1–13.

Lima-Melo Y, Gollan PJ, Tikkanen M, Silveira JA, Aro E. 2019. Consequences of photosystem-I damage and repair on photosynthesis and carbon use in *Arabidopsis thaliana*. The Plant Journal 97, 1061–1072.

Luu Trinh MD, Miyazaki D, Ono S, et al. 2021. The evolutionary conserved iron-sulfur protein TCR controls P700 oxidation in photosystem I. iScience 24, 102059.

Maekawa S, Ohnishi M, Wada S, Ifuku K, Miyake C. 2024. Enhanced reduction of ferredoxin in PGR5-deficient mutant of *Arabidopsis thaliana* stimulated ferredoxin-dependent cyclic electron flow around photosystem I. International Journal of Molecular Sciences 25, 2677.

Martín M, Casano LM, Zapata JM, Guéra A, Del Campo EM, Schmitz-Linneweber C, Maier RM, Sabater B. 2004. Role of thylakoid Ndh complex and peroxidase in the protection against photo-oxidative stress: fluorescence and enzyme activities in wild-type and *ndhF*-deficient tobacco. Physiologia Plantarum 122, 443–452.

Messant M, Hani U, Lai T, Wilson A, Shimakawa G, Krieger-Liszkay A. 2024. Plastid terminal oxidase (PTOX) protects photosystem I and not photosystem II against photoinhibition in *Arabidopsis thaliana* and *Marchantia polymorpha*. The Plant Journal 117, 669–678.

Miller NT, Vaughn MD, Burnap RL. 2021. Electron flow through NDH-1 complexes is the major driver of cyclic electron flow-dependent proton pumping in cyanobacteria. Biochimica et Biophysica Acta (BBA)-Bioenergetics 1862, 148354.

Miyake C. 2020. Molecular mechanism of oxidation of P700 and suppression of ROS production in photosystem I in response to electron-sink limitations in C_3_ plants. Antioxidants 9, 230.

Nakamura A, Ogawa T, Shimakawa G, Kobayashi R, Shikanai T, Munekage YN. 2026. Cyclic electron flow around photosystem I provides energy production and photoprotection in C_4_ plants. Plant Physiology. 10.1093/plphys/kiag146.

Nakamura Y, Wada S, Miyake C, Makino A, Suzuki Y. 2024. Regulation of photosystems II and I depending on N partitioning to Rubisco in rice leaves: a study using Rubisco-antisense transgenic plants. Journal of Plant Research 137, 1165–1175.

Napaumpaiporn P, Ogawa T, Sonoike K, Nishiyama Y. 2024. Improved capacity for the repair of photosystem II via reinforcement of the translational and antioxidation systems in *Synechocystis* sp. PCC 6803. The Plant Journal 117, 1165–1178.

Nashilevitz S, Melamed-Bessudo C, Izkovich Y, Rogachev I, Osorio S, Itkin M, Adato A, Pankratov I, Hirschberg J, Fernie AR. 2010. An orange ripening mutant links plastid NAD(P)H dehydrogenase complex activity to central and specialized metabolism during tomato fruit maturation. The Plant Cell 22, 1977–1997.

Nelson N, Junge W. 2015. Structure and energy transfer in photosystems of oxygenic photosynthesis. Annual Review of Biochemistry 84, 659–683.

Nikkanen L, Santana Sanchez A, Ermakova M, Rögner M, Cournac L, Allahverdiyeva Y. 2020. Functional redundancy between flavodiiron proteins and NDH-1 in *Synechocystis* sp. PCC 6803. The Plant Journal 103, 1460–1476.

Nikkanen L, Toivola J, Trotta A, Diaz MG, Tikkanen M, Aro E, Rintamäki E. 2018. Regulation of cyclic electron flow by chloroplast NADPH-dependent thioredoxin system. Plant Direct 2, 1–24.

Nishiyama Y, Yamamoto H, Allakhverdiev SI, Inaba M, Yokota A, Murata N. 2001. Oxidative stress inhibits the repair of photodamage to the photosynthetic machinery. The EMBO journal 20, 5587–5594.

Obara A, Ogawa M, Oyama Y, Suzuki Y, Kono M. 2022. Effects of high irradiance and low water temperature on photoinhibition and repair of photosystems in Marimo (*Aegagropila linnaei*) in Lake Akan, Japan. International Journal of Molecular Sciences 24, 60.

Okegawa Y, Motohashi K. 2020. M-type thioredoxins regulate the PGR5/PGRL1-dependent pathway by forming a disulfide-linked complex with PGRL1. The Plant Cell 32, 3866–3883.

Opatíková M, Kouřil R. 2024. Unique structural attributes of the PSI-NDH supercomplex in *Physcomitrium patens*. The Plant Journal 120, 2226–2237.

Otani T, Kato Y, Shikanai T. 2018. Specific substitutions of light-harvesting complex I proteins associated with photosystem I are required for supercomplex formation with chloroplast NADH dehydrogenase-like complex. The Plant Journal 94, 122–130.

Otani T, Yamamoto H, Shikanai T. 2017. Stromal loop of Lhca6 is responsible for the linker function required for the NDH–PSI supercomplex formation. Plant and Cell Physiology 58, 851–861.

Ozaki H, Mizokami Y, Sugiura D, Sohtome T, Miyake C, Sakai H, Noguchi K. 2023. Tight relationship between two photosystems is robust in rice leaves under various nitrogen conditions. Journal of Plant Research 136, 201–210.

Ozawa S-I, Zhang G, Sakamoto W. 2024. Dysfunction of chloroplast protease activity mitigates *pgr5* phenotype in the green algae *Chlamydomonas reinhardtii*. Plants 13, 606.

Pan X, Cao D, Xie F, Xu F, Su X, Mi H, Zhang X, Li M. 2020. Structural basis for electron transport mechanism of complex I-like photosynthetic NAD(P)H dehydrogenase. Nature Communications 11, 610.

Peeler TC, Naylor AW. 1988. A comparison of the effects of chilling on thylakoid electron transfer in pea (*Pisum sativum* L.) and cucumber (*Cucumis sativus* L.). Plant Physiology 86, 147–151.

Rajagopal S, Joly D, Gauthier A, Beauregard M, Carpentier R. 2005. Protective effect of active oxygen scavengers on protein degradation and photochemical function in photosystem I submembrane fractions during light stress. The FEBS Journal 272, 892–902.

Richardson KH, Wright JJ, Šimėnas M, Thiemann J, Esteves AM, McGuire G, Myers WK, Morton JJ, Hippler M, Nowaczyk MM. 2021. Functional basis of electron transport within photosynthetic complex I. Nature Communications 12, 5387.

Rodriguez-Heredia M, Saccon F, Wilson S, Finazzi G, Ruban AV, Hanke GT. 2022. Protection of photosystem I during sudden light stress depends on ferredoxin: NADP(H) reductase abundance and interactions. Plant Physiology 188, 1028–1042.

Rutherford AW, Osyczka A, Rappaport F. 2012. Back-reactions, short-circuits, leaks and other energy wasteful reactions in biological electron transfer: redox tuning to survive life in O_2_. FEBS Letters 586, 603–616.

Satoh H, Ohara Y, Hanke G, Ifuku K, Shimakawa G, Suzuki Y, Makino A, Moriguchi K, Miyake C. 2025. The regulation of PSI cyclic electron transport by both plastoquinone and ferredoxin redox states: correlation with the rate of proton motive force utilization. Frontiers in Plant Science, 16, 1626163.

Schuller JM, Birrell JA, Tanaka H, Konuma T, Wulfhorst H, Cox N, Schuller SK, Thiemann J, Lubitz W, Setif P. 2019. Structural adaptations of photosynthetic complex I enable ferredoxin-dependent electron transfer. Science 363, 257–260.

Shaku K, Shimakawa G, Hashiguchi M, Miyake C. 2016. Reduction-induced suppression of electron flow (RISE) in the photosynthetic electron transport system of *Synechococcus elongatus* PCC 7942. Plant and Cell Physiology 57, 1443–1453.

Shen L, Tang K, Wang W, Wang C, Wu H, Mao Z, An S, Chang S, Kuang T, Shen J-R. 2022. Architecture of the chloroplast PSI–NDH supercomplex in *Hordeum vulgare*. Nature 601, 649–654.

Shikanai T. 2016. Chloroplast NDH: a different enzyme with a structure similar to that of respiratory NADH dehydrogenase. Biochimica et Biophysica Acta (BBA)-Bioenergetics 1857, 1015–1022.

Shikanai T. 2024. Molecular genetic dissection of the regulatory network of proton motive force in chloroplasts. Plant and Cell Physiology 65, 537–550.

Shikanai T, Endo T, Hashimoto T, Yamada Y, Asada K, Yokota A. 1998. Directed disruption of the tobacco *ndhB* gene impairs cyclic electron flow around photosystem I. Proceedings of the National Academy of Sciences 95, 9705–9709.

Shikanai T, Ieda H, Kobayashi Y, Tamura MN. 2025. The chloroplast NADH dehydrogenase-like complex: evolutionary considerations. Plant and Cell Physiology 66, 1525–1535.

Shimakawa G, Miyake C. 2018. Oxidation of P700 ensures robust photosynthesis. Frontiers in Plant Science 9, 1617.

Shimakawa G, Müller P, Miyake C, Krieger-Liszkay A, Sétif P. 2024. Photo-oxidative damage of photosystem I by repetitive flashes and chilling stress in cucumber leaves. Biochimica et Biophysica Acta (BBA)-Bioenergetics 1685, 149490.

Shimakawa G, Murakami A, Niwa K, Matsuda Y, Wada A, Miyake C. 2019. Comparative analysis of strategies to prepare electron sinks in aquatic photoautotrophs. Photosynthesis Research 139, 401–411.

Shuvalov V, Nuijs A, Van Gorkom H, Smit H, Duysens L. 1986. Picosecond absorbance changes upon selective excitation of the primary electron donor P-700 in photosystem I. Biochimica et Biophysica Acta (BBA)-Bioenergetics 850, 319–323.

Sonoike K. 1999. The different roles of chilling temperatures in the photoinhibition of photosystem I and photosystem II. Journal of Photochemistry and Photobiology B: Biology 48, 136–141.

Sonoike K, Terashima I. 1994. Mechanism of photosystem-I photoinhibition in leaves of *Cucumis sativus* L.. Planta 194, 287–293.

Storti M, Puggioni MP, Segalla A, Morosinotto T, Alboresi A. 2020. The chloroplast NADH dehydrogenase-like complex influences the photosynthetic activity of the moss *Physcomitrella patens*. Journal of Experimental Botany 71, 5538–5548.

Strand DD, Fisher N, Kramer DM. 2017. The higher plant plastid NAD(P)H dehydrogenase-like complex (NDH) is a high efficiency proton pump that increases ATP production by cyclic electron flow. Journal of Biological Chemistry 292, 11850–11860.

Su X, Cao D, Pan X, Shi L, Liu Z, Dall’Osto L, Bassi R, Zhang X, Li M. 2022. Supramolecular assembly of chloroplast NADH dehydrogenase-like complex with photosystem I from *Arabidopsis thaliana*. Molecular Plant 15, 454–467.

Suganami M, Konno S, Maruhashi R, et al. 2022. Expression of flavodiiron protein rescues defects in electron transport around PSI resulting from overproduction of Rubisco activase in rice. Journal of Experimental Botany 73, 2589–2600.

Sun Y, Geng Q, Du Y, Yang X, Zhai H. 2017. Induction of cyclic electron flow around photosystem I during heat stress in grape leaves. Plant Science 256, 65–71.

Sun H, Shi Q, Liu N-Y, Zhang S-B, Huang W. 2023. Drought stress delays photosynthetic induction and accelerates photoinhibition under short-term fluctuating light in tomato. Plant Physiology and Biochemistry 196, 152–161.

Takagi D, Amako K, Hashiguchi M, et al. 2017*a*. Chloroplastic ATP synthase builds up a proton motive force preventing production of reactive oxygen species in photosystem I. The Plant Journal 91, 306–324.

Takagi D, Ihara H, Takumi S, Miyake C. 2019. Growth light environment changes the sensitivity of photosystem I photoinhibition depending on common wheat cultivars. Frontiers in Plant Science 10, 686.

Takagi D, Ishizaki K, Hanawa H, Mabuchi T, Shimakawa G, Yamamoto H, Miyake C. 2017b. Diversity of strategies for escaping reactive oxygen species production within photosystem I among land plants: P700 oxidation system is prerequisite for alleviating photoinhibition in photosystem I. Physiologia Plantarum 161, 56–74.

Takagi D, Kishie A, Ifuku K. 2026. Photosystem I photoinhibition attenuates LHCI-dependent light-harvesting activity in spinach leaves. Plant Physiology kiag251, 10.1093/plphys/kiag251.

Takagi D, Tani S. 2023. Impact of growth light environment on oxygen sensitivity in rice: Pseudo-first-order response of photosystem I photoinhibition to O_2_ partial pressure. Physiologia Plantarum 175, e14009.

Takeuchi K, Che Y, Nakano T, Miyake C, Ifuku K. 2022. The ability of P700 oxidation in photosystem I reflects chilling stress tolerance in cucumber. Journal of Plant Research 135, 681–692.

Takeuchi K, Harimoto S, Che Y, Kumazawa M, Satoh H, Maekawa S, Miyake C, Ifuku K. 2025*a*. The protective role of chloroplast NADH dehydrogenase-like complex (NDH) against PSI photoinhibition under chilling stress. New Phytologist 248, 2262–2279.

Takeuchi K, Harimoto S, Maekawa S, Miyake C, Ifuku K. 2025*b*. PSII photoinhibition as a protective strategy: maintaining an oxidative state of PSI by suppressing PSII activity under environmental stress. Physiologia plantarum 177, e70392.

Takeuchi K, Ochiai K, Kobayashi M, Kuroda K, Ifuku K. 2024. Light-chilling stress causes hyper-accumulation of iron in shoot, exacerbating leaf oxidative damage in cucumber. Plant and Cell Physiology 65, 1873–1887.

Terashima I, Funayama S, Sonoike K. 1994. The site of photoinhibition in leaves of *Cucumis sativus* L. at low temperatures is photosystem I, not photosystem II. Planta 193, 300–306.

Terashima I, Huang L-K, Osmond CB. 1989. Effects of leaf chilling on thylakoid functions, measured at room temperature, in *Cucumis sativus* L. and *Oryza sativa* L. Plant and Cell Physiology 30, 841–850.

Terashima I, Kashino Y, Katoh S. 1991*a*. Exposure of leaves of *Cucumis sativus* L. to low temperatures in the light causes uncoupling of thylakoids I. Studies with isolated thylakoids. Plant and Cell Physiology 32, 1267–1274.

Terashima I, Matsuo M, Suzuki Y, Yamori W, Kono M. 2021. Photosystem I in low light-grown leaves of *Alocasia odora*, a shade-tolerant plant, is resistant to fluctuating light-induced photoinhibition. Photosynthesis Research 149, 69–82.

Terashima I, Sonoike K, Kawazu T, Katoh S. 1991*b*. Exposure of leaves of *Cucumis sativus* L. to low temperatures in the light causes uncoupling of thylakoids II. Non-destructive measurements with intact leaves. Plant and Cell Physiology 32, 1275–1283.

Tikhonov AN. 2024. The cytochrome *b*_6_*f* complex: Plastoquinol oxidation and regulation of electron transport in chloroplasts. Photosynthesis Research 159, 203–227.

Tikkanen M, Mekala NR, Aro E-M. 2014. Photosystem II photoinhibition-repair cycle protects Photosystem I from irreversible damage. Biochimica et Biophysica Acta (BBA)-Bioenergetics 1837, 210–215.

Tiwari A, Mamedov F, Fitzpatrick D, Gunell S, Tikkanen M, Aro E-M. 2024. Differential FeS cluster photodamage plays a critical role in regulating excess electron flow through photosystem I. Nature Plants 10, 1592–1603.

Tyystjärvi E. 2008. Photoinhibition of photosystem II and photodamage of the oxygen evolving manganese cluster. Coordination Chemistry Reviews 252, 361–376.

Ueda M, Kuniyoshi T, Yamamoto H, Sugimoto K, Ishizaki K, Kohchi T, Nishimura Y, Shikanai T. 2012. Composition and physiological function of the chloroplast NADH dehydrogenase-like complex in *Marchantia polymorpha*. The Plant Journal 72, 683–693.

Urzinger S, Avramova V, Frey M, Urbany C, Scheuermann D, Presterl T, Reuscher S, Ernst K, Mayer M, Marcon C. 2025. Embracing native diversity to enhance the maximum quantum efficiency of photosystem II in maize. Plant Physiology 197, kiae670.

Vicino P, Carrillo J, Gómez R, Shahinnia F, Tula S, Melzer M, Rutten T, Carrillo N, Hajirezaei M-R, Lodeyro AF. 2021. Expression of flavodiiron proteins Flv2-Flv4 in chloroplasts of Arabidopsis and tobacco plants provides multiple stress tolerance. International Journal of Molecular Sciences 22, 1178.

Wada S, Takagi D, Miyake C, Makino A, Suzuki Y. 2019. Responses of the photosynthetic electron transport reactions stimulate the oxidation of the reaction center chlorophyll of photosystem I, P700, under drought and high temperatures in rice. International journal of Molecular Sciences 20, 2068.

Wada S, Yamamoto H, Suzuki Y, Yamori W, Shikanai T, Makino A. 2018. Flavodiiron protein substitutes for cyclic electron flow without competing CO_2_ assimilation in rice. Plant Physiology 176, 1509–1518.

Wakasugi T, Tsudzuki J, Ito S, Nakashima K, Tsudzuki T, Sugiura M. 1994. Loss of all *ndh* genes as determined by sequencing the entire chloroplast genome of the black pine *Pinus thunbergii*. Proceedings of the National Academy of Sciences 91, 9794–9798.

Wang P, Duan W, Takabayashi A, Endo T, Shikanai T, Ye J-Y, Mi H. 2006. Chloroplastic NAD(P)H dehydrogenase in tobacco leaves functions in alleviation of oxidative damage caused by temperature stress. Plant Physiology 141, 465–474.

Wang Y, Luo A, Lyu M-JA, Wang Y-M, Huang Y, Ni X, Yang J, Wen Y, Zhu X-G. 2026. Inducibility of cyclic electron transport is linked to the transition from C_3_–C_4_ to C_4_ photosynthesis in *Flaveria*. *Plant Physiology*, kiag089.

Wang F, Yan J, Ahammed GJ, Wang X, Bu X, Xiang H, Li Y, Lu J, Liu Y, Qi H. 2020. PGR5/PGRL1 and NDH mediate far-red light-induced photoprotection in response to chilling stress in tomato. Frontiers in Plant Science 11, 669.

Wu X, Shu S, Wang Y, Yuan R, Guo S. 2019. Exogenous putrescine alleviates photoinhibition caused by salt stress through cooperation with cyclic electron flow in cucumber. Photosynthesis Research 141, 303–314.

Yadav KS, Semchonok DA, Nosek L, Kouřil R, Fucile G, Boekema EJ, Eichacker LA. 2017. Supercomplexes of plant photosystem I with cytochrome *b*_6_*f*, light-harvesting complex II and NDH. Biochimica et Biophysica Acta (BBA)-Bioenergetics 1858, 12–20.

Yamamoto H, Sato N, Shikanai T. 2021. Critical role of NdhA in the incorporation of the peripheral arm into the membrane-embedded part of the chloroplast NADH dehydrogenase-like complex. Plant and Cell Physiology 62, 1131–1145.

Yamamoto H, Takahashi S, Badger MR, Shikanai T. 2016. Artificial remodelling of alternative electron flow by flavodiiron proteins in *Arabidopsis*. Nature Plants 2, 1–7.

Yamori W, Sakata N, Suzuki Y, Shikanai T, Makino A. 2011. Cyclic electron flow around photosystem I via chloroplast NAD(P)H dehydrogenase (NDH) complex performs a significant physiological role during photosynthesis and plant growth at low temperature in rice. The Plant Journal 68, 966–976.

Zhang Y, Fan Y, Lv X, Zeng X, Zhang Q, Wang P. 2023. Deficiency in NDH-cyclic electron transport retards heat acclimation of photosynthesis in tobacco over day and night shift. Frontiers in Plant Science 14, 1267191.

Zhang Y, He S, Chen G. 2024. Metabolomics of chloroplasts combined with photosynthetic properties reveal low-temperature stress responses in sugarcane (*Saccharum officinarum* L.). Journal of Plant Growth Regulation 44, 748–764.

Zhang Z-S, Jin L-Q, Li Y-T, Tikkanen M, Li Q-M, Ai X-Z, Gao H-Y. 2016. Ultraviolet-B radiation (UV-B) relieves chilling-light-induced PSI photoinhibition and accelerates the recovery of CO_2_ assimilation in cucumber (*Cucumis sativus* L.) leaves. Scientific Reports 6, 34455.

Zhang Q, Tian S, Chen G, Tang Q, Zhang Y, Fleming AJ, Zhu X, Wang P. 2024. Regulatory NADH dehydrogenase-like complex optimizes C_4_ photosynthetic carbon flow and cellular redox in maize. New Phytologist 241, 82–101.

Zhang Z, Zhang M, Burnap RL. 2025. Action at a distance: The remarkable coupling of CO_2_ uptake to electron transfer in specialized cyanobacterial NDH-1 complexes. Proceedings of the National Academy of Sciences 122, e2511786122.

Zheng M, Jiang Y, Ran Z, Liang S, Xiao T, Li X, Ma W. 2025. A cyanobacteria-derived intermolecular salt bridge stabilizes photosynthetic NDH-1 and prevents oxidative stress. Communications Biology 8, 172.

Zhou Y, Huang L, Zhang Y, Shi K, Yu J, Nogués S. 2007. Chill-induced decrease in capacity of RuBP carboxylation and associated H_2_O_2_ accumulation in cucumber leaves are alleviated by grafting onto figleaf gourd. Annals of Botany 100, 839–848.

Zhou Q, Yamamoto H, Shikanai T. 2023. Distinct contribution of two cyclic electron transport pathways to P700 oxidation. Plant Physiology 192, 326–341.

Zivcak M, Brestic M, Kunderlikova K, Sytar O, Allakhverdiev SI. 2015. Repetitive light pulse-induced photoinhibition of photosystem I severely affects CO_2_ assimilation and photoprotection in wheat leaves. Photosynthesis Research 126, 449–463.

