## Supplementary Fig. for "The PSI–NDH supercomplex prevents chilling-induced PSI photoinhibition"

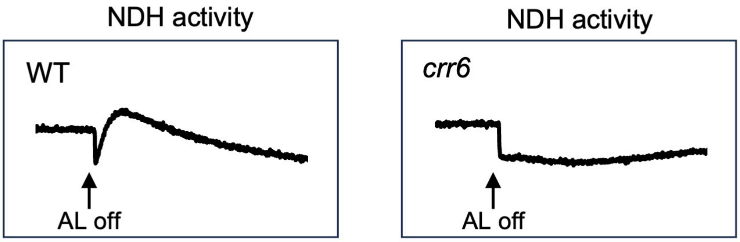


**Supplementary Fig. S1.** Post-illumination fluorescence rise (PIFR) analyzed under ambient conditions (23 ºC) in WT (Hitomebore) and *crr6*. Leaves were kept in the dark for at least 20 min, followed by exposure to AL for 5 min. The subsequent transient increase in Chl fluorescence after switching off AL was monitored in the dark.

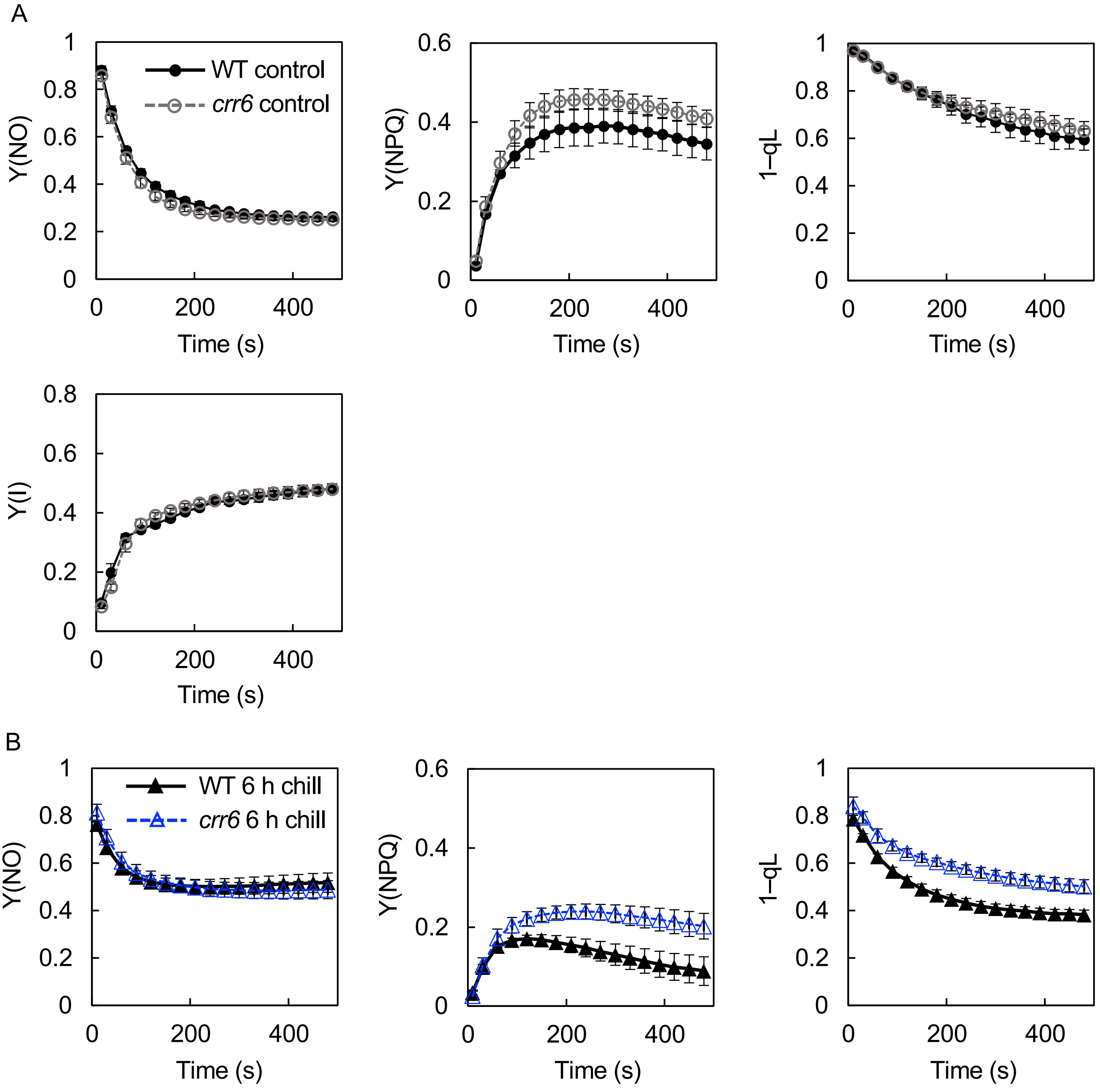


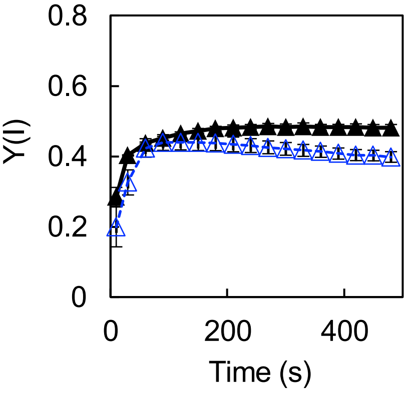


**Supplementary Fig. S2.** Photosynthetic activity before and after chilling stress in WT (Hitomebore) and *crr6*. Experimental conditions and leaves were identical to those in Fig. 2A–F. **A**. Before chilling stress (Control). **B**. After 6 h of chilling stress. Y(NO) represents the quantum yield of nonregulated energy dissipation in PSII, Y(NPQ) represents the quantum yield of regulated non-photochemical quenching, qL represents the fraction of open PSII reaction centers based on the lake model, and Y(I) represents the fraction of P700 that can be oxidized by SP.

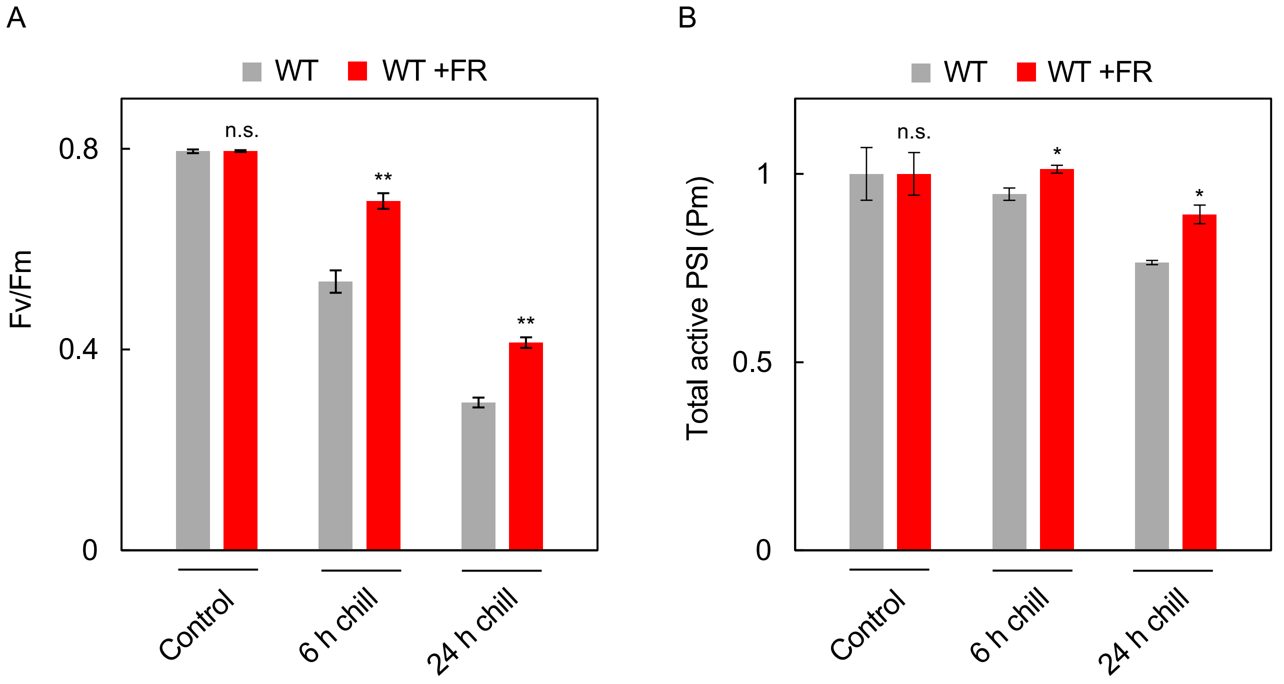


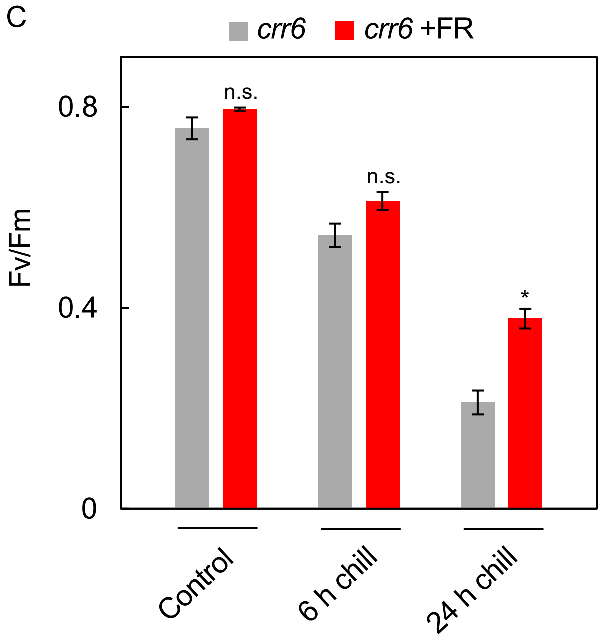


**Supplementary Fig. S3.** Protective effects of FR light on PSI and PSII. Experimental conditions were identical to those shown in Fig. 2I. **A**. Fv/Fm measured before and after chilling stress in WT in the presence or absence of FR light. **B**. Pm measured before and after chilling stress in WT in the presence or absence of FR light. **C**. Fv/Fm measured before and after chilling stress in *crr6* in the presence or absence of FR light. Values are the mean ± SE, n = 6, biological replicates. Asterisks (*, **) indicate statistically significant differences ($p$<0.01 and $p$<0.001, respectively), whereas 'n.s.' denotes no significant difference ($p$>0.05) (Student's $t$-test).

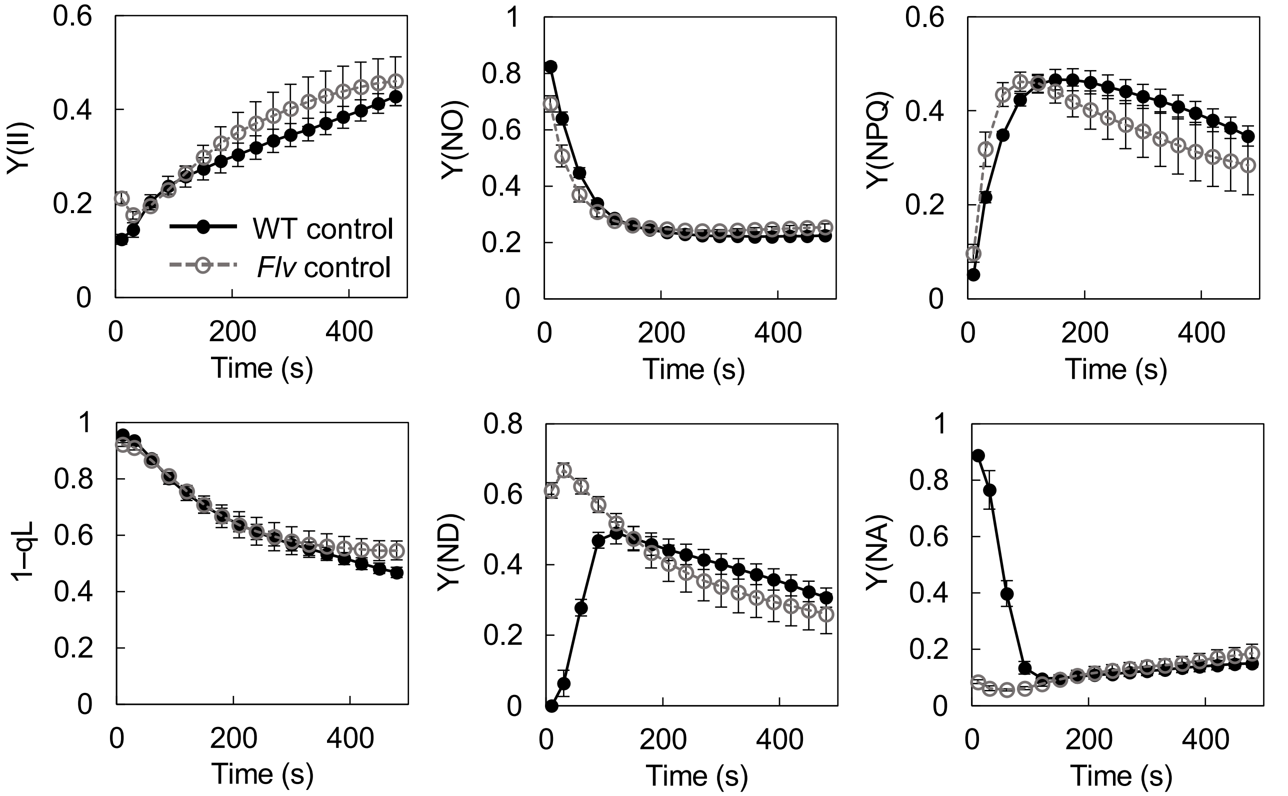

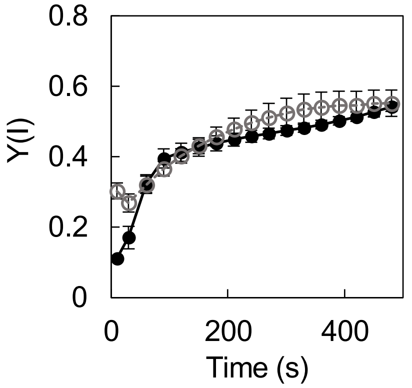


**Supplementary Fig. S4.** Photosynthetic activity before chilling stress (Control) in WT (Nipponbare) and *Flv*-expressing rice. Experimental conditions were identical to those in Fig. 3F–H, except that chilling stress was not applied.

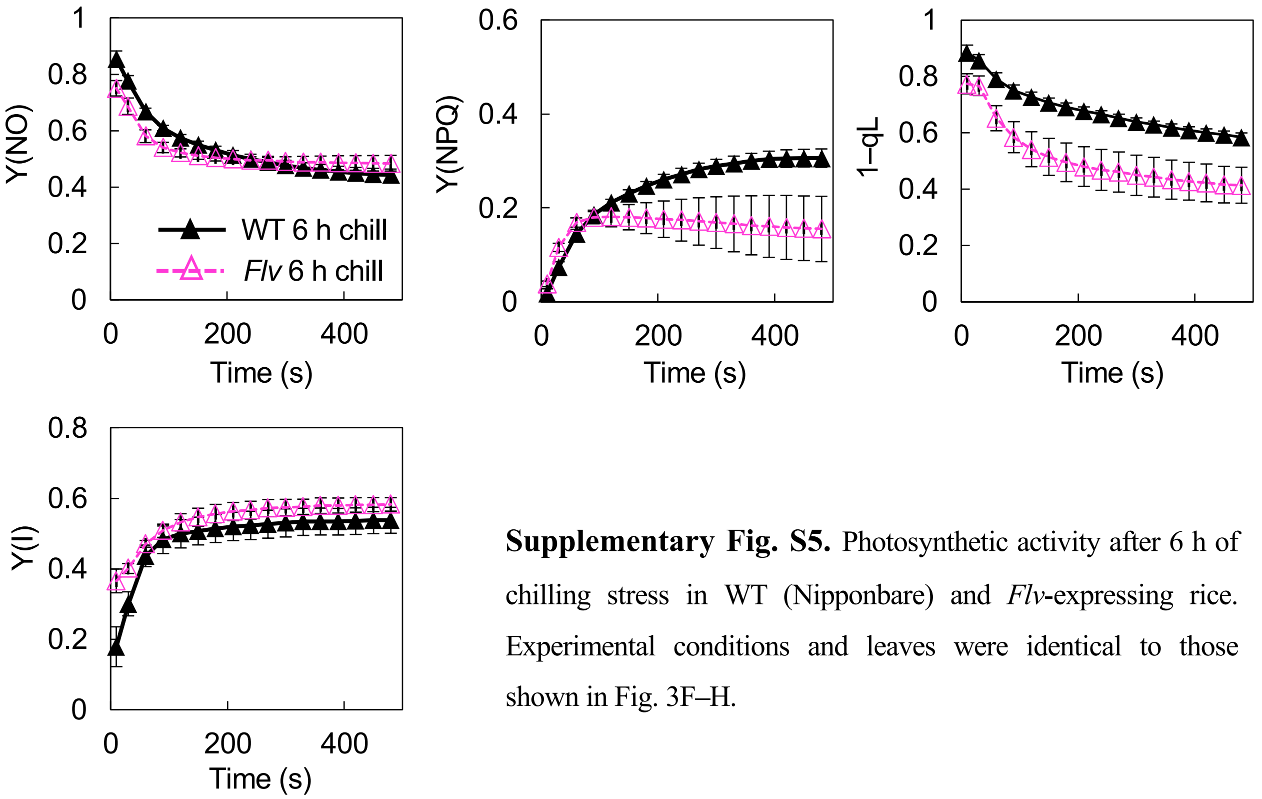


**Supplementary Fig. S5.** Photosynthetic activity after 6 h of chilling stress in WT (Nipponbare) and *Flv*-expressing rice. Experimental conditions and leaves were identical to those shown in Fig. 3F–H.

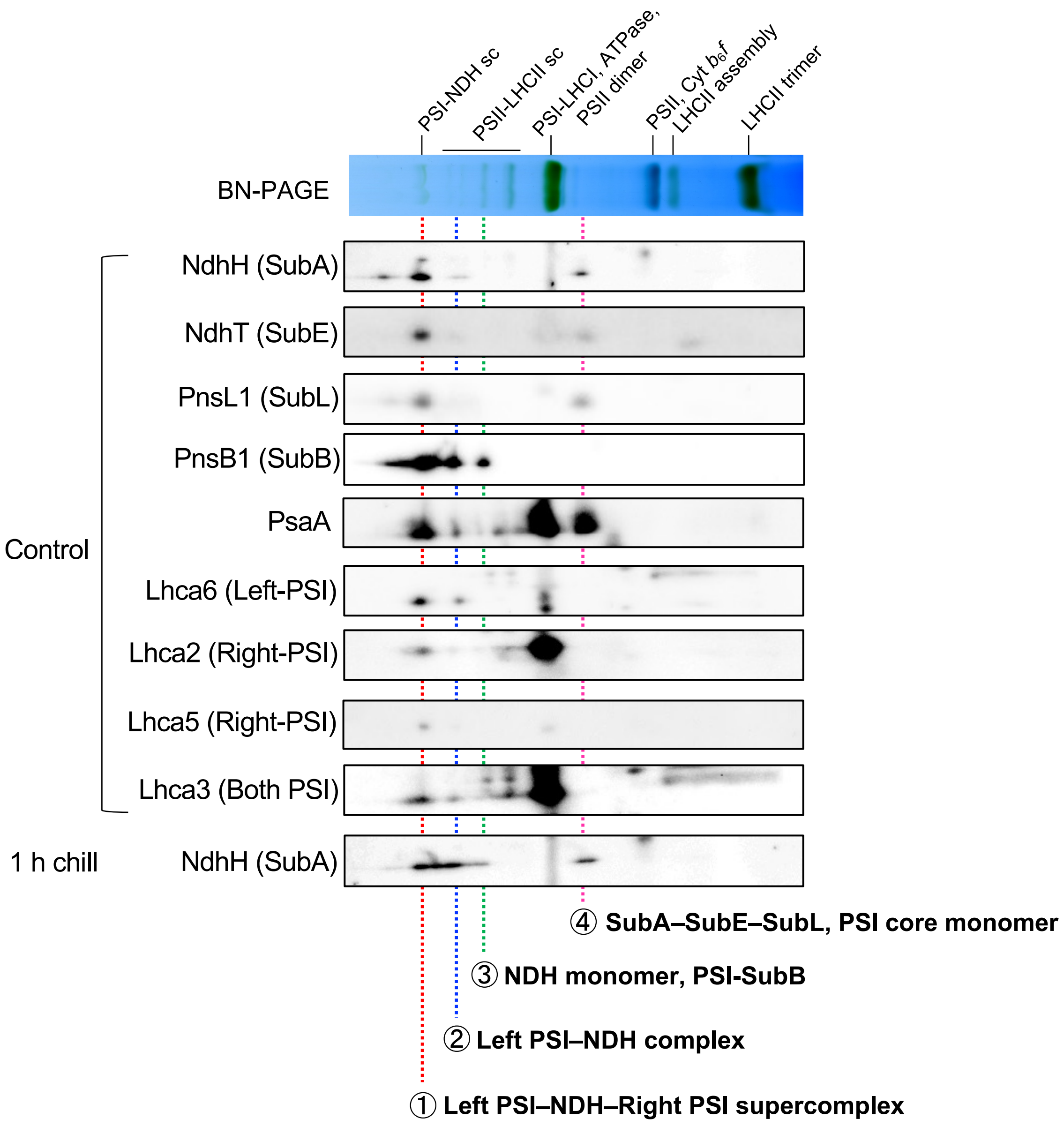


**Supplementary Fig. S6.** Analysis of the PSI–NDH sc assembly state in cucumber. Thylakoid membrane proteins isolated before and after 1 h of chilling stress were separated by BN-PAGE and subsequently analyzed by 2D-SDS-PAGE. NDH subunits (NdhH, NdhT, PnsL1, and PnsB1) and PSI–LHCI subunits (PsaA, Lhca2, Lhca3, Lhca5, and Lhca6) were detected using specific antibodies. The experimental procedure was identical to that shown in Fig. 5, and part of the data includes results presented in Fig. 5 and previously reported data (Takeuchi *et al*., 2025, *New Phytologist*). Red dotted line **1** indicates the PSI–NDH–PSI sc band; blue dotted line **2** indicates the left PSI–NDH complex band; green dotted line **3** indicates the band position corresponding to the NDH monomer and the PSI–SubB assembly intermediate; and magenta dotted line **4** indicates the band corresponding to the SubA–SubE–SubL assembly intermediate and the PSI core monomer.

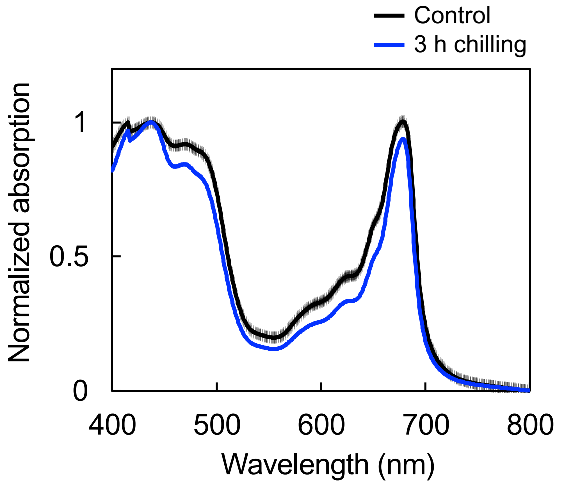


**Supplementary Fig. S7.** Absorption spectra of isolated thylakoids before and after chilling stress in cucumber. Isolated thylakoids (50 μg Chl $mL^{-1}$) obtained before and after 3 h of chilling stress were suspended in HEPES buffer. Absorption spectra (400–800 nm) were recorded at 25 ºC using a UV-2600 spectrophotometer (Shimadzu, Kyoto, Japan) and normalized to the absorption peak at 430 nm. Values are the mean ± SE, n = 3–4.

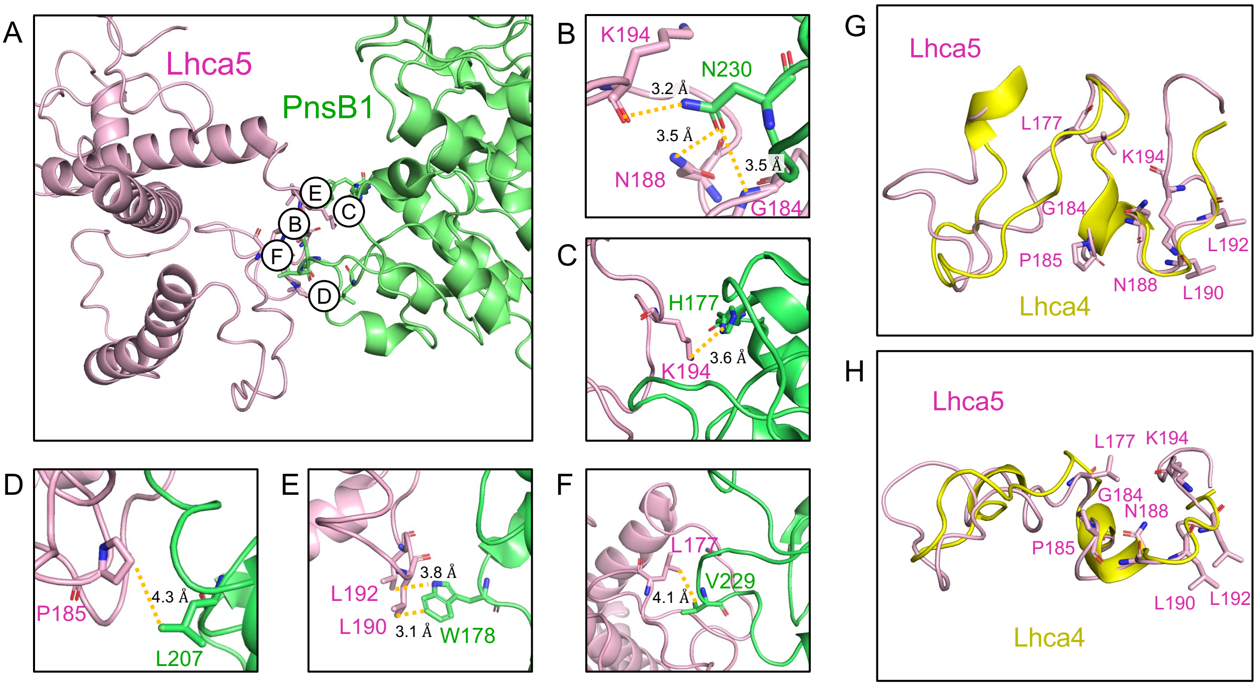


**Supplementary Fig. S8**. Interaction sites between Lhca5 and PnsB1. Because the amino acid sequences and overall structures of Lhca5 and Lhca4 are highly similar, NDH binding modes were inferred by identifying residues and structural features specific to Lhca5. Analysis of the PSI–NDH sc cryo-EM structure (PDB ID: 7WG5) revealed an interaction between the stromal A–C loop region of Lhca5 and PnsB1 (Kato *et al.*, 2018*a*; Su *et al.*, 2022). Structural inspection using PyMOL (version 3.1.0) suggested that the interaction involves hydrogen bonds between Lhca5 G184, N188, or the main chain of K194 and PnsB1, a cation–π interaction involving the side chain of K194, and three hydrophobic interactions involving P185, L190/L192, and L177. Among these residues, interspecies comparison of Lhca4 and Lhca5 sequences retrieved from the NCBI database identified P185, L192, and K194 as Lhca5-specific residues. P185 and L192 were completely conserved within Lhca5, whereas K194 was conserved as a basic residue (Lys or Arg) (Supplementary Fig. S10). Structural superposition of Lhca4 and Lhca5 using PyMOL showed that the overall conformation of the stromal loop region was largely conserved; however, in Lhca5, the loop position of the K194 main chain was closer to PnsB1 N230 than in Lhca4, supporting the formation of a hydrogen bond between the K194 main chain of Lhca5 and PnsB1 N230. **A**. Overall structure of the Lhca5-PnsB1 interacting site (pink: Lhca5, light green: PnsB1). **B–F**. Magnified views of regions B–F in panel A, showing detailed interaction sites. **B**. Hydrogen-bonding sites involving the main chain of K194, N188 or G184. **C**. Cation–π interaction site involving the side chain of K194. **D**. Hydrophobic interaction site involving P185. **E**. Hydrophobic interaction sites involving L190 and L192. **F**. Hydrophobic interaction site involving L177. **G–H**. Structural superposition of Lhca4 and Lhca5 at the stromal loop regions. Both structures were obtained from the PSI–NDH sc cryo-EM structure (PDB ID: 7WG5).

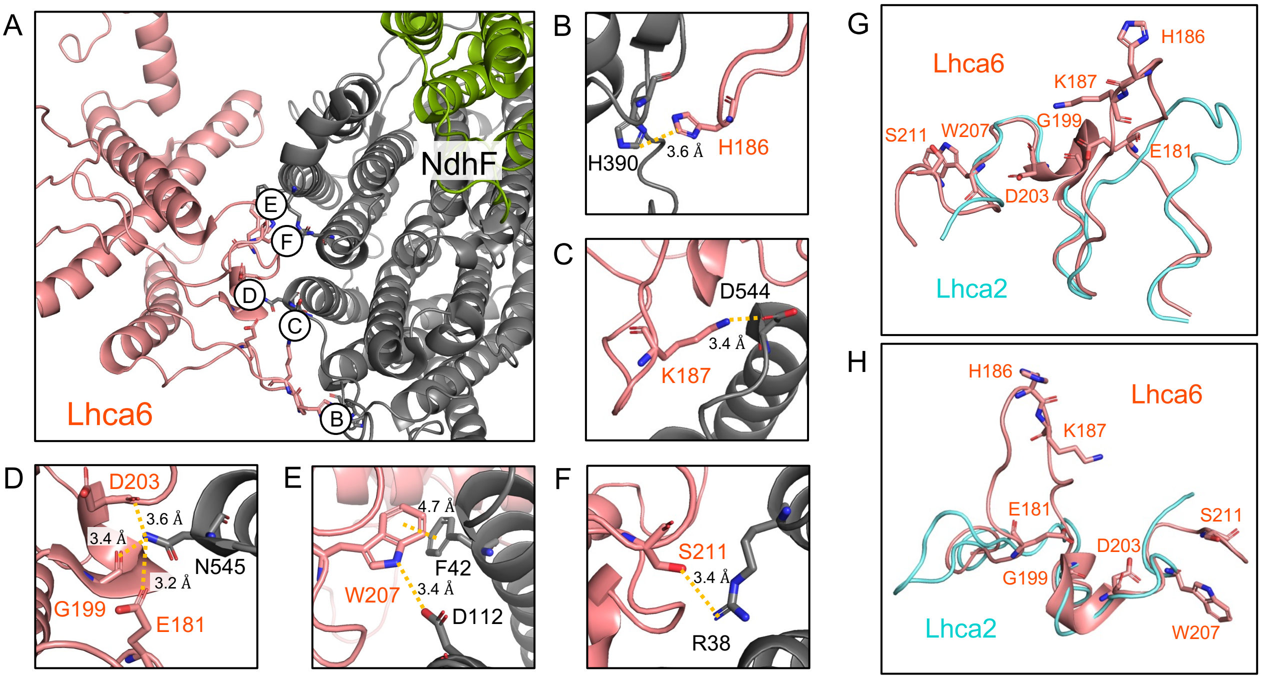


**Supplementary Fig. S9.** Interaction sites between Lhca6 and NdhF. Because the amino acid sequences and overall structures of Lhca6 and Lhca2 are highly similar, NDH binding modes were inferred by identifying residues and structural features specific to Lhca6. Using a similar approach as Supplementary Fig. S8, an interaction between the stromal A–C loop region of Lhca6 and NdhF was identified (Otani *et al.*, 2017; Su *et al.*, 2022; Introini *et al.*, 2025). Structural analysis suggested the involvement of hydrogen bonds formed by E181, G199, or D203, as well as hydrogen bonds involving H186 and S211, a salt bridge involving K187, and a π–π interaction involving W207. Comparison of Lhca2 and Lhca6 sequences identified E181 and K187 as Lhca6-specific residues (Supplementary Fig. S11). E181 was conserved in all but three species, and K187 was conserved as a basic residue (Lys or Arg) in most species except for some leguminous plants. In *Spinacia oleracea*, where Arg is present at position 187, hydrogen bonding at this site has been suggested (Introini *et al.*, 2025). Structural comparison between Lhca2 and Lhca6 further revealed differences at positions E181, H186, K187, and S211. **A**. Overall structure of Lhca6-NdhF interacting site (orange: Lhca6, black: NdhF). **B–F**. Magnified views of regions B–F in panel A, showing detailed interaction sites. **B**. Hydrogen-bonding site involving H186. **C**. Salt bridge site involving K187. **D**. Hydrogen-bonding sites involving D203, G199, or E181. **E**. π–π interaction site involving W207. **F**. Hydrogen-bonding site involving S211. **G–H**. Structural superposition of Lhca2 and Lhca6 at the stromal loop regions. Both structures were obtained from PSI-NDH sc structure PDB ID: 7WG5.

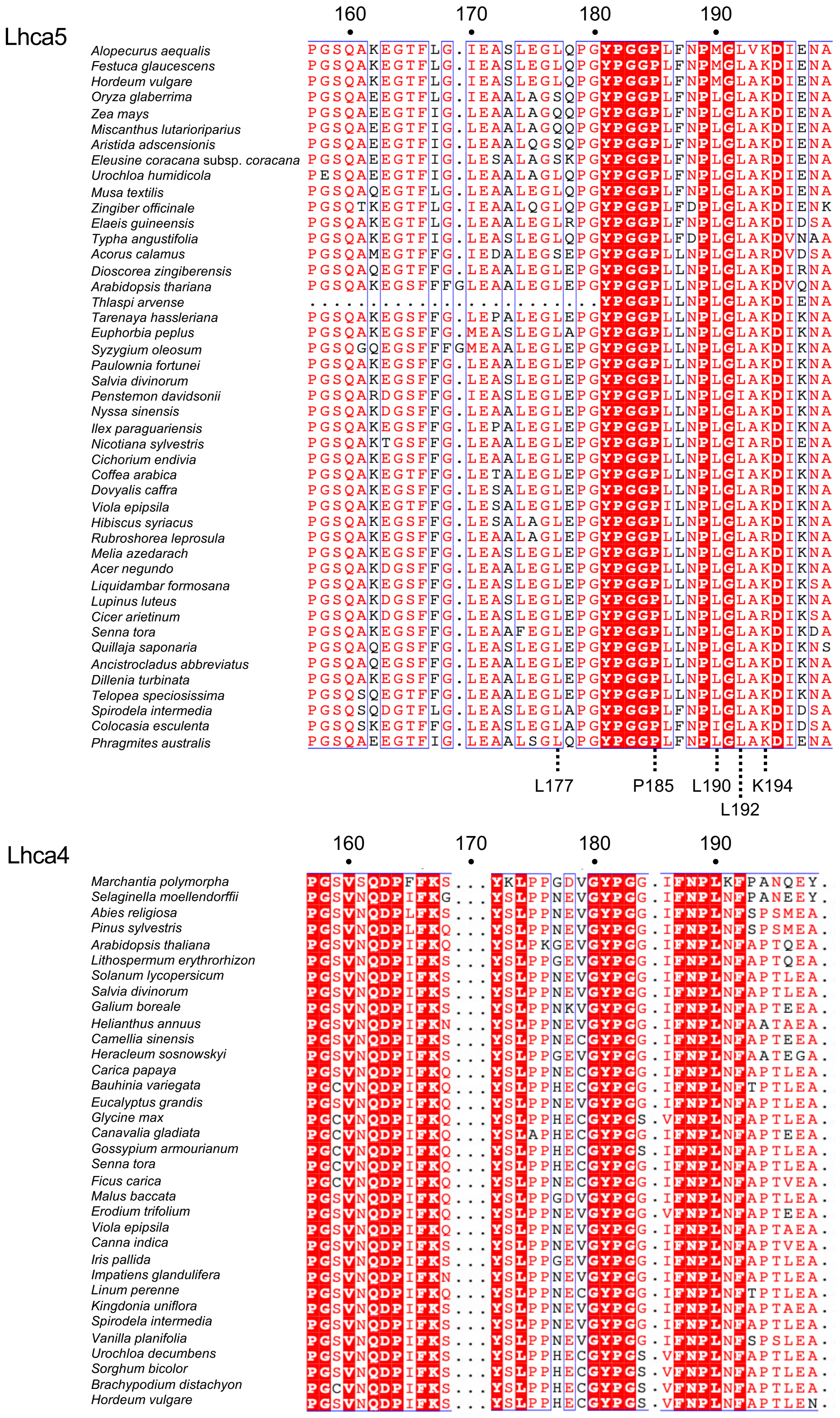


**Supplementary Fig. S10.** Comparison of amino acid sequences in the stromal loop regions of Lhca4 and Lhca5. Lhca4 and Lhca5 sequences were retrieved by BLASTP searches using *Arabidopsis thaliana* Lhca4 and Lhca5 as queries, yielding 34 Lhca4 and 45 Lhca5 sequences from different plant species. Multiple sequence alignments were generated using ClustalW. Residues that are identical across all sequences are highlighted with a red background, whereas conserved residues with similar physicochemical properties are shown in red text. Highly conserved sequence motifs shared among all sequences are further indicated by blue boxes.

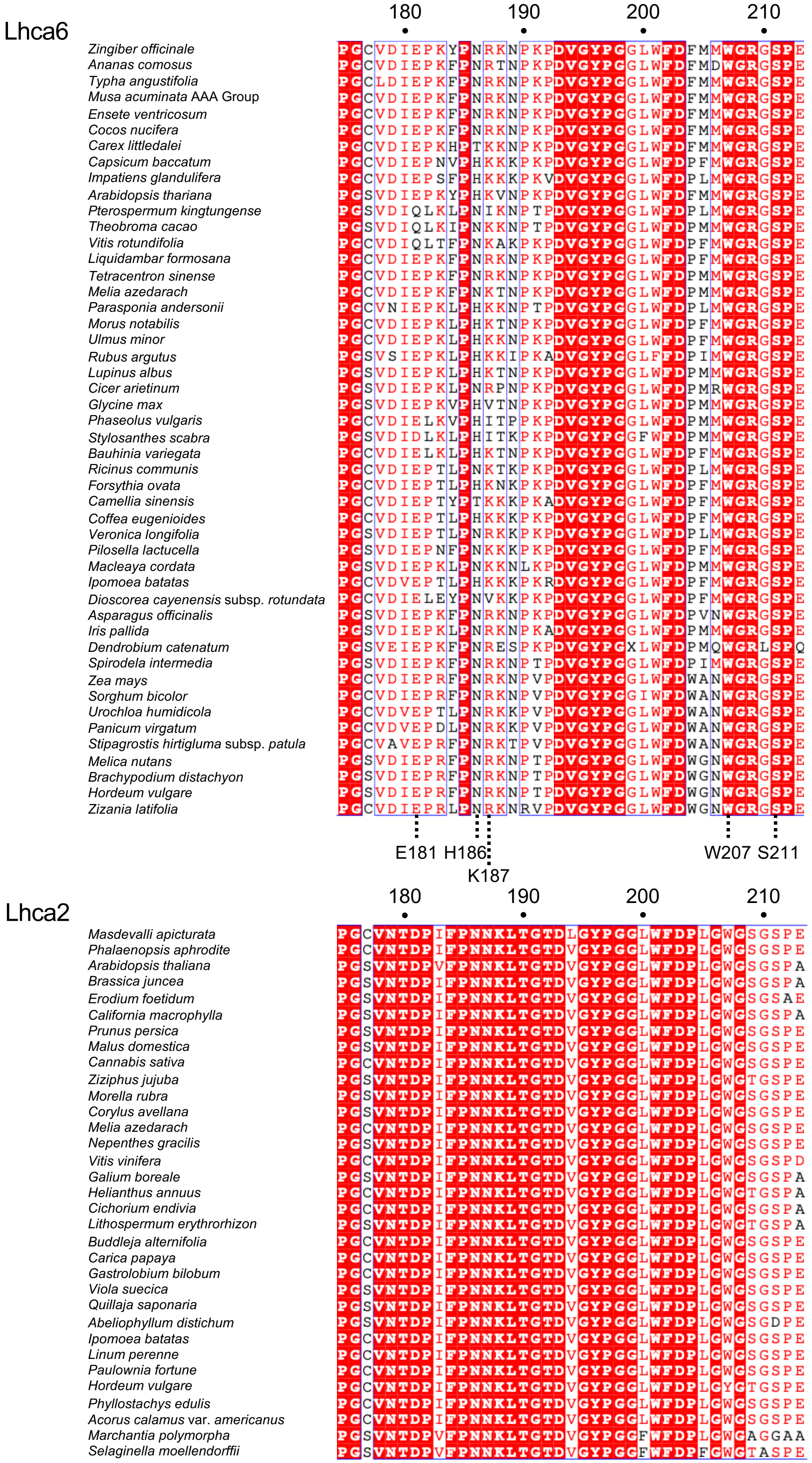


**Supplementary Fig. S11.** Comparison of amino acid sequences in the stromal loop regions of Lhca2 and Lhca6. Lhca2 and Lhca6 sequences were retrieved by BLASTP searches using Arabidopsis thaliana Lhca2 and Lhca6 as query sequences, yielding 33 Lhca2 and 48 Lhca6 sequences from different plant species. Multiple sequence alignments were generated using ClustalW. Identical residues across all sequences are highlighted with a red background, whereas conserved residues with similar physicochemical properties are shown in red text. Highly conserved sequence motifs shared among all sequences are further indicated by blue boxes.
